# Apathy–anhedonia is associated with pessimistic beliefs yet sharpened goal-directed planning

**DOI:** 10.1101/2025.08.14.670362

**Authors:** Shiyi Liang, Evan M. Russek, Robb B. Rutledge, G. Elliott Wimmer

## Abstract

A core premise in decision-making research is that beliefs about uncertain outcomes shape value-based choice: pessimistic expectations should reduce goal pursuit, whereas optimistic expectations should promote it. Depression provides a critical test of this assumption, as motivational symptoms such as apathy and anhedonia are linked to diminished expectations of success. Yet whether such biases actually impair goal-directed behavior, particularly when risk must be learned from experience, remains unknown. We developed a multi-step planning task in which participants (n = 384) learned the structure and risks of maze-like environments across two days, then chose between certain rewards and risky goals varying in reward magnitude and distance, with greater distance entailing higher compounding risk. Using transdiagnostic symptom factors, we observed a dissociation: individuals with elevated apathy–anhedonia reported lower subjective expectations of goal success, yet showed no reduction in risky goal pursuit. Instead, higher apathy–anhedonia was associated with enhanced performance, including stronger discrimination of goal values, increased model-based flexibility, and faster goal navigation. These effects were specific to apathy–anhedonia; other symptom factors, including worry and impulsivity, were unrelated to expectations or goal-directed behavior. Together, these findings demonstrate that pessimistic beliefs and goal-directed competence are dissociable, revealing that motivational symptoms can coexist with sharpened value-based planning when risk is learned from experience.

## Introduction

A common assumption in decision science is that subjective beliefs about uncertain outcomes guide value-based choice (Rangel et al., 2008; Tversky and Kahneman, 1992). On this view, pessimistic expectations should reduce goal pursuit and optimistic expectations should promote it. Depression provides a critical test of this assumption, because motivational symptoms such as apathy and anhedonia are linked to diminished expectations of success and reduced reward anticipation (Husain and Roiser, 2018; Strunk and Adler, 2009). Yet whether such pessimistic beliefs actually constrain goal-directed behavior remains unclear.

Existing research has focused primarily on how psychiatric symptoms relate to the quality of goal-directed, or model-based, control. This leaves two more basic questions unresolved: whether motivational symptoms reduce pursuit of uncertain goals, as distinct from willingness to expend effort for reward (Treadway et al., 2012), and whether explicit expectations and choice behavior correspond within the same learned environment.

Regarding model-based control, the picture is mixed. Large transdiagnostic studies using the two-step task have consistently linked compulsivity and intrusive thought symptom factors to reduced model-based control, but have found no such relationship for depression-related symptoms (Donegan et al., 2023; Gillan et al., 2016; Patzelt et al., 2019). Other studies, including some in clinical samples, have reported mixed results ranging from impairment to intact performance (Gillan et al., 2020; Heller et al., 2018; Heo et al., 2021), and most recently, a large-scale study reported enhanced model-based control at higher levels of depression symptoms (Sookud et al., 2025). One limitation, however, is that apathy–anhedonia is rarely isolated from broader depression and anxiety dimensions, leaving its specific relationship with goal-directed behavior unclear.

The picture is further complicated by the tasks themselves. Real-world goal pursuit typically depends on learning environmental structure through repeated experience, drawing on consolidated memory, and evaluating risk that compounds across sequential steps. In such settings, altered expectations about future outcomes could plausibly decrease or reduce the quality of goal pursuit (Korn et al., 2014; Strunk and Adler, 2009). Yet most laboratory paradigms probe goal-directed control under relatively shallow planning horizons and simplified uncertainty, while lacking measures of expectations. Paradigms that combine multi-step planning with learned uncertainty may therefore be necessary to test how motivational symptoms shape both what people expect and what they choose (Scholl and Klein-Flügge, 2018; Wise et al., 2024).

We developed a paradigm that places these demands at the center of the task. Participants (n = 384) learned the structure and risks of maze-like environments through extended experience across two days (**Figure 1**). During a subsequent choice phase, participants decided between certain rewards and risky goals varying in reward magnitude and distance, with greater distance entailing higher compounded risk of failure. Crucially, participants also provided explicit estimates of goal success, enabling a direct comparison between beliefs and choices. Transdiagnostic symptom factors derived from clinical questionnaires isolated motivational symptoms (apathy–anhedonia) from worry and impulsivity.

**Figure 1.**
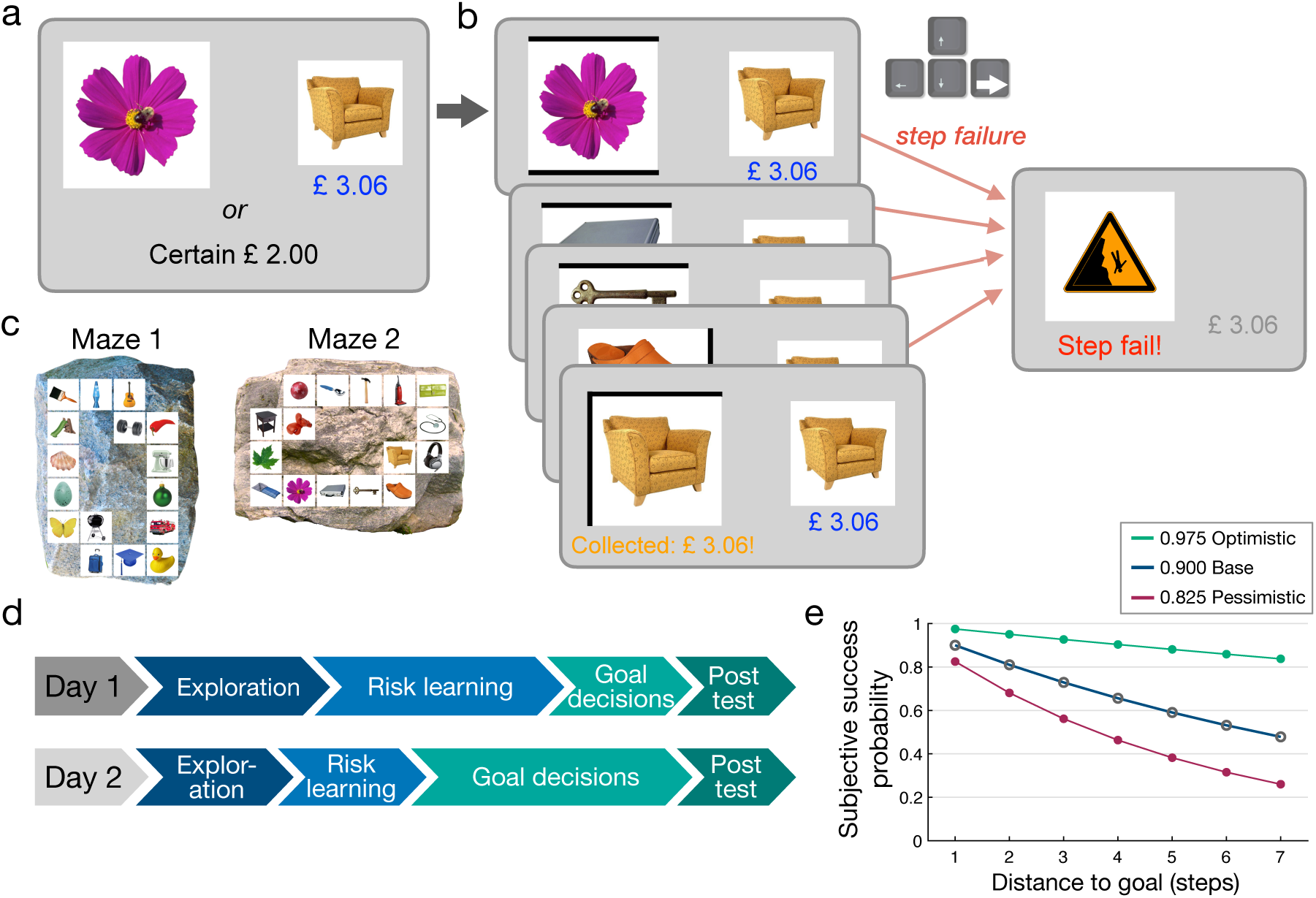
Risky goal task. a) Choice phase. Participants chose between a certain alternative and a risky goal option, defined by a start location, goal location, and reward. In this example shown, the start is the flower and the goal is the chair, four steps away via the shorter path, yielding £3.06 if reached (see Maze 2 in panel c). b) Goal pursuit. Shown during learning trials and after choosing a goal. Participants selected a navigation direction at each step using the arrow keys. Black lines indicate walls; encountering a wall prompted reselection. Each step carried an undisclosed ∼10% probability of failure, independent of participant actions (pink arrows), which terminated the trial (outcome screen, right; see also **Figure S1**). c) Maze layouts. Each maze contained 16 locations marked by unique objects arranged in a loop. This overhead view was shown briefly between task phases but was not available during trials. d) Experimental timeline. Each day included exploration, learning, and choice phases. Subjective success ratings and location memory assessments followed each session, with additional memory tests early in each day. A follow-up session (day 3 or 4) included success ratings and location memory only. e) Expected success probability illustration. Success probability as a function of subjective step failure rate. For a goal 6 steps away, expected success ranges from 86% under an optimistic estimate to 53% at baseline and 32% under a pessimistic estimate.

We identified a dissociation between beliefs and behavior in individuals with elevated apathy–anhedonia symptoms. As predicted, higher apathy–anhedonia was associated with more pessimistic expectations of goal success. However, these pessimistic beliefs did not translate into reduced goal pursuit. Instead, higher apathy–anhedonia was associated with enhanced goal-directed performance across convergent measures, including increased sensitivity to goal value, more flexible adjustment to changing contingencies, and faster goal-directed navigation. Together, these findings challenge the view that motivational symptoms reflect a generalized deficit in goal-directed control, demonstrating that pessimistic explicit expectations can diverge from value-guided planning when uncertainty is learned from experience.

## Results

### Risky goal task

We examined goal-directed decision-making under learned uncertainty and its relationship with transdiagnostic psychiatric symptoms. The study was preregistered (https://osf.io/x3z5f); throughout, we use the term ‘hypothesized’ to denote preregistered hypotheses.

The risky goal task models a common real-world problem: deciding whether a potentially valuable goal is worth pursuing when the probability of success must be estimated from past experience and declines with the number of steps required to reach it. Across two days, participants learned and navigated two loop-shaped “mazes,” in which each location was represented by a unique image (**Figure 1c**). On goal pursuit trials, participants started from a designated location and attempted to reach a goal by selecting either the clockwise or counter-clockwise path (**Figure 1b**; **Figure S1**).

Critically, each step carried a small, independent probability failure (∼10%) that terminated the trial. This probability was not disclosed but had to be learned through experience. Because failure risk compounded across steps, the expected value of goal pursuit decreased with distance – longer paths could yield greater rewards but also carried greater cumulative risk of failure (**Figure 1e; Figure S2**).

To quantify how participants valued risky goals, the task included a choice phase in which participants decided between a certain reward and a goal option defined by a start item, goal item, and reward value (**Figure 1a**). If participants chose the goal, they navigated toward it by selecting an initial direction and making subsequent movement choices at each step; if they chose the certain alternative, that outcome was shown followed by a matched delay. Goal distance varied from 1–7 steps (defined by the shorter path), and offered reward values generally increased with distance. Reward values were adjusted adaptively across trials to estimate each participant’s indifference point at each distance – the reward at which they were equally likely to choose the risky goal or the certain option.

Several additional task components probed distinct aspects of goal-directed behavior. Occasional “roadblock” trials tested flexible use of the learned model by temporarily blocking a signaled location (**Figure S3**). Participants also provided subjective ratings of goal success probability for a separate set of start-goal combinations, allowing us to measure explicit beliefs within the learned environment. Finally, memory for object locations was assessed early and late in each session to quantify structure knowledge independent of decision-making.

The experiment was conducted across two days to support extended learning of both of environmental structure and step-wise risk, more closely approximating how such knowledge is acquired outside the laboratory. Each day included exploration, learning, and choice phases, with a longer choice phase on day two. Primary analyses focus on the day two choice phase, after participants had more fully learned the environments.

### Transdiagnostic symptom dimensions

To examine how psychiatric symptoms related to task behavior, we adopted a transdiagnostic approach that captures symptom dimensions cutting across diagnostic categories (Gillan et al., 2016; Wise et al., 2023). Factor analysis of responses to a broad set of clinical questionnaires identified three factors, labeled ‘Apathy–Anhedonia’, ‘Worry’, and ‘Impulsivity’ based on item loadings (**Figure 2**; **Figure S4-S5**; **Supp. Results**). To better differentiate depression from anxiety symptoms, our set included additional depression- and anxiety-focused scales beyond those used in earlier factor-analytic work (Gillan et al., 2016; Wise et al., 2023). Participants also completed two general risk propensity questionnaires analyzed separately (Frey et al., 2017)(**Supplemental Results**).

**Figure 2.**
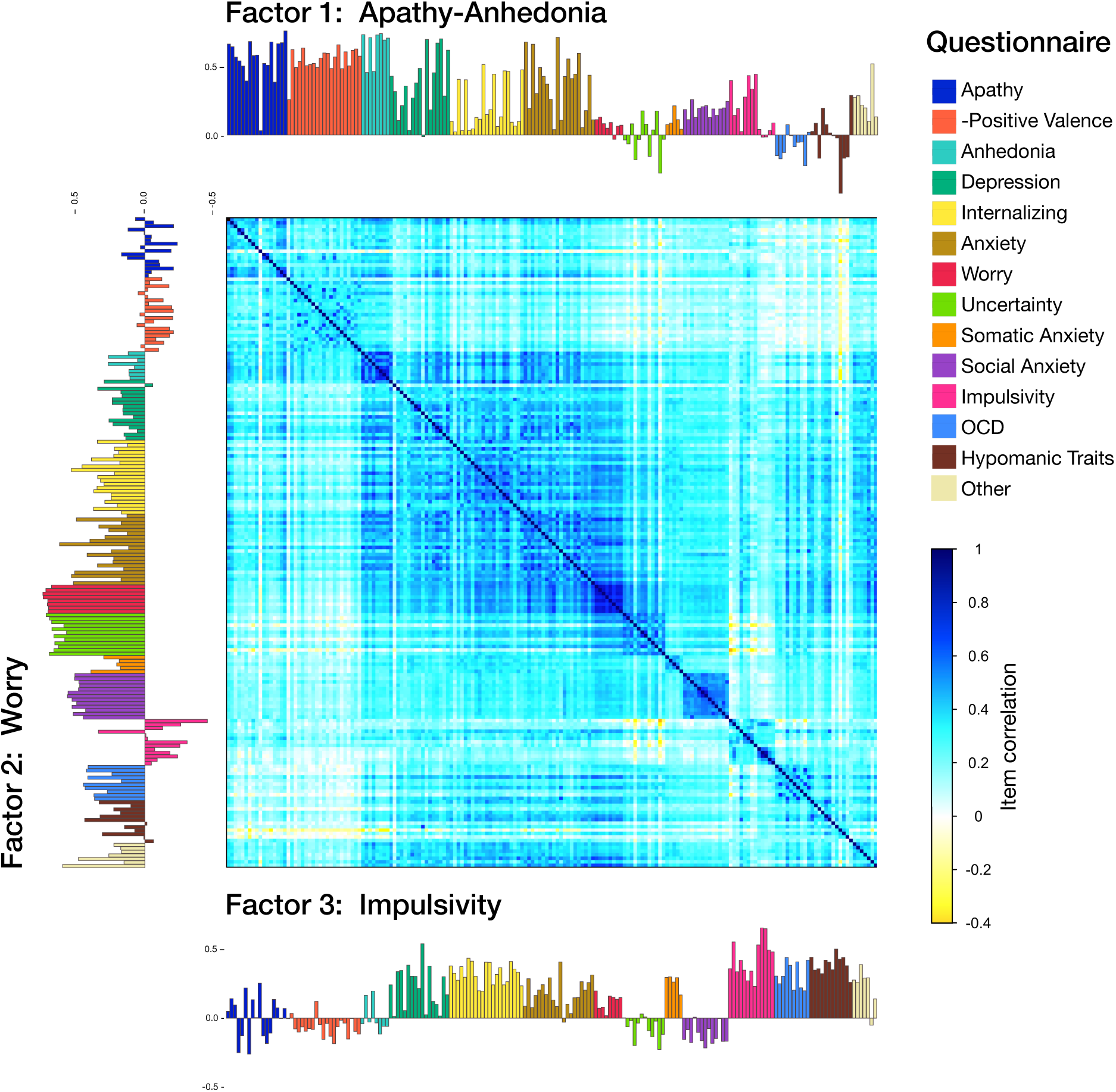
Transdiagnostic symptom dimensions. Center: correlation matrix of the 184 questionnaire items entered into the factor analysis. Three factors were identified, labeled ‘Apathy–Anhedonia’, ‘Worry’, and ‘Impulsivity’ based on item loadings. Item loadings displayed along the top, left and bottom edges of the matrix; loading bar colors indicate the source questionnaire for each item. Items from the positive valence scale were sign-inverted for visualization. See Methods for full questionnaire names; ‘Other’ refers to depression-targeted items drawn from additional scales.

### Learning and memory

Participants completed location memory assessments early and late on each day, in which they replaced individual items onto an empty maze grid (**Figure S1b**). Location memory was already high following the initial exploration phase on day one (83.89 ± 13.92% [SD]; **Figure 3a**), indicating that participants rapidly built internal models of the two environments. First-move direction accuracy during goal pursuit, defined as pursuing the shortest path, was 86.45 ± 16.42% on day one, confirming a strong model-based component to navigation. Note that poor location memory and first-move accuracy were used as exclusion criteria to ensure analyses focused on participants who had successfully learned the environments (**Supp. Methods**).

**Figure 3.**
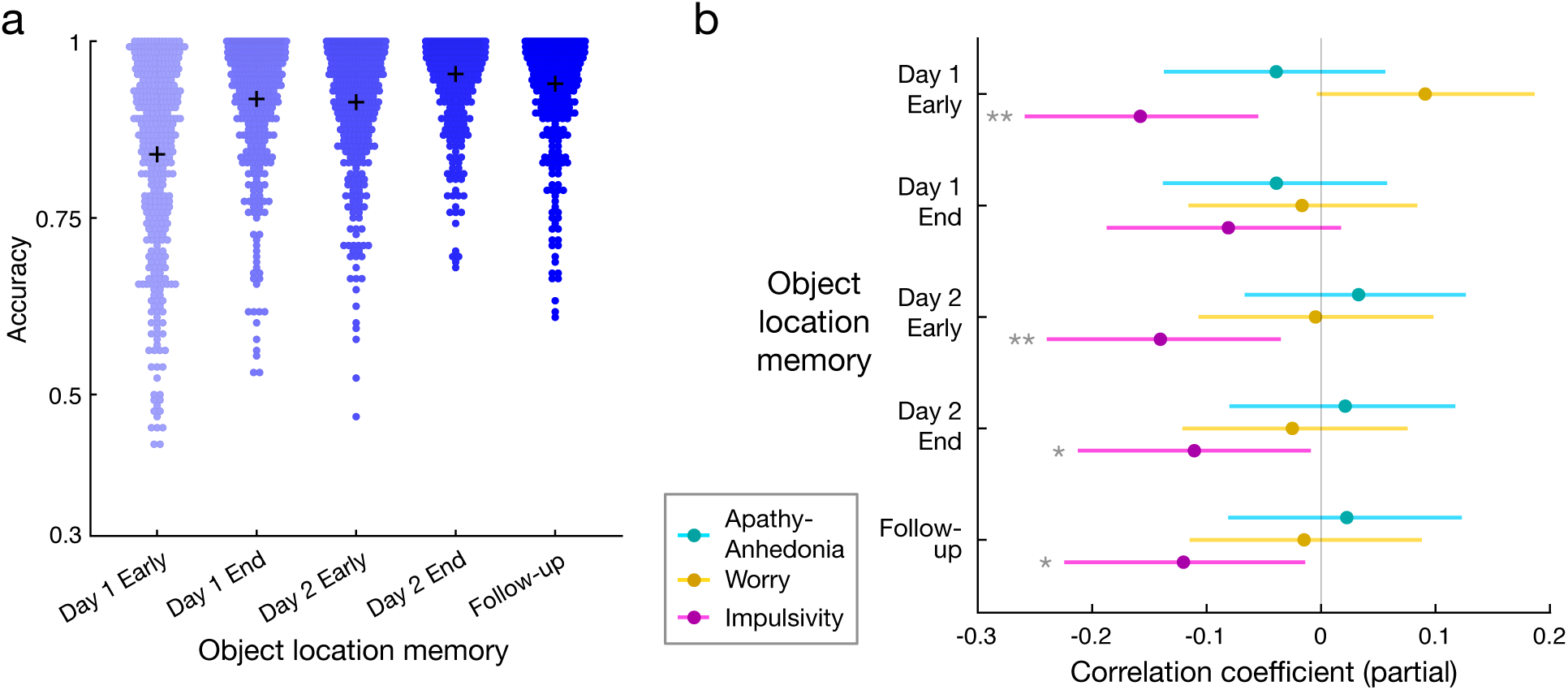
Memory and transdiagnostic factors. a) Performance in object location replacement memory tests across time. Points represent individual participants, with black crosses indicating the mean. b) Location memory performance correlation with factor scores. All analyses include the three transdiagnostic factors as simultaneous predictors. Confidence intervals (lines) were estimated by bootstrapping. * p < 0.05; ** p < 0.01.

We tested the preregistered hypothesis that symptoms related to compulsivity and intrusive thoughts (CIT) would be associated with weaker acquisition of model-based knowledge of the maze (Seow et al., 2021; Sharp et al., 2023; Sookud et al., 2025). Of the three factors, ‘Impulsivity,’ which also heavily weighted intrusive thought items, most closely corresponded to the previously described CIT dimension. Consistent with this prediction, higher ‘Impulsivity’ scores were associated with poorer location memory across multiple assessments, including both memory tests from the primary day two session (early; partial r_s_(380) = −0.141, p = 0.0060; late: partial r_s_(380) = −0.111, p = 0.0304; partial Spearman correlations controlling for the other factors; **Figure 3b**), the early day one test (early: partial r_s_(381) = −0.158, p = 0.0020), and the day three follow-up (partial r_s_(368) = −0.120, p = 0.0206), though the late day one assessment was not significant (partial r_s_(380) = −0.080, p = 0.114). Bivariate analyses showed the same pattern (**Figure S6**). Neither ‘Worry’ nor ‘Apathy–Anhedonia’ scores were significantly related to location memory (p-values > 0.161; **Figure 3**), contrary to our hypothesis that depression-related symptoms might also relate to poorer memory (Dillon and Pizzagalli, 2018).

### Goal success ratings

To measure explicit beliefs about future success in the learned environments, participants provided subjective predictions of goal success probability for specific start-goal combinations varying in distance (**Figure S7**; distance information was not explicitly provided). As expected, these ratings decreased strongly with distance (t = - 31.619, p < 0.001, multilevel model; **Figure S7**), indicating that participants tracked the increasing risk associated with longer paths. On average, ratings were close to but slightly more pessimistic than experienced success rates (mean rating: 63.58 ± 0.60%; extrapolated experienced success rate: 65.49 ± 0.24%; t_(389)_ = −3.0389, p = 0.0025).

Ratings were also highly stable across sessions, with a strong correlation between day two and a follow-up at least one day later (r(371) = 0.810, p < 0.001), indicating that these estimates reflected stable beliefs.

We hypothesized that depression-related motivational symptoms would be associated with lower expectations of goal success. Supporting this prediction, higher ‘Apathy–Anhedonia’ factor scores were associated with lower success ratings (t_(369)_ = −2.756, p = 0.006; regression controlling for other factor scores; **Figure 4**). This relationship was present earlier in learning on day one (t_(377)_ = −2.082, p = 0.0380) and persisted in the follow-up session (t_(365)_ = −2.633; p = 0.009), suggesting a stable pessimistic bias rather than a transient effect. Neither ‘Worry’ nor ‘Impulsivity’ scores were significantly associated with success ratings (p-values > 0.33, uncorrected; **Figure 4**; **Figure S9**).

**Figure 4.**
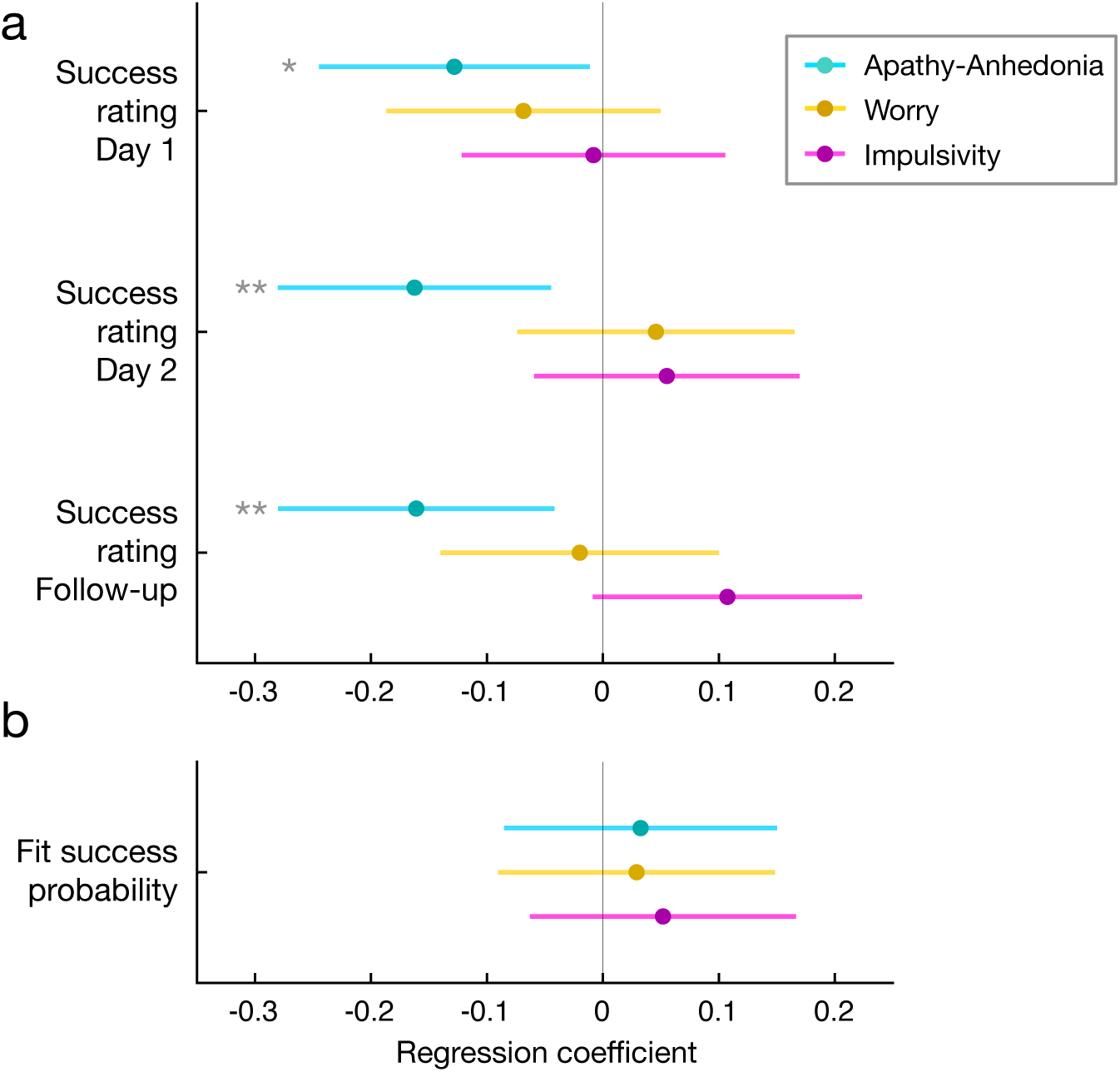
Dissociation between subjective beliefs and choice behavior in apathy–anhedonia. a) Higher ‘Apathy–Anhedonia’ scores were associated with lower subjective goal success ratings, reflecting pessimistic expectations. b) In contrast, ‘Apathy–Anhedonia’ scores were unrelated to model-estimated step success probability (risk-taking for goals). All analyses include the three transdiagnostic factors as simultaneous predictors. * p < 0.05; ** p < 0.01.

The ‘Apathy–Anhedonia’ effect remained significant when controlling for location memory performance, memory confidence, experienced step success rate, or baseline mood. This is notable because baseline mood itself was strongly negatively correlated with ‘Apathy–Anhedonia’ scores (**Figure S10**). These results indicate that motivational symptoms are associated with more pessimistic beliefs about goal success, and that this effect cannot be explained by differences in learning, experience, or general affect.

### Risky goal decisions and computational model

Having established that Apathy–Anhedonia is associated with pessimistic success expectations, we next asked whether these beliefs were accompanied by reduced goal pursuit. During the choice phase, participants decided between a certain reward and risky goals varying in reward magnitude and distance. As expected, participants were more likely to choose the goal option as its reward increased (z = 14.76, p < 0.001, multilevel model with fixed effects of participant and distance), and they required larger rewards to pursue more distant goals (indifference points increasing with distance: t = 20.496, p < 0.001, multilevel model; **Figure 5**). This pattern indicates that participants integrated knowledge of the environment with learned step-wise risk when making decisions.

**Figure 5.**
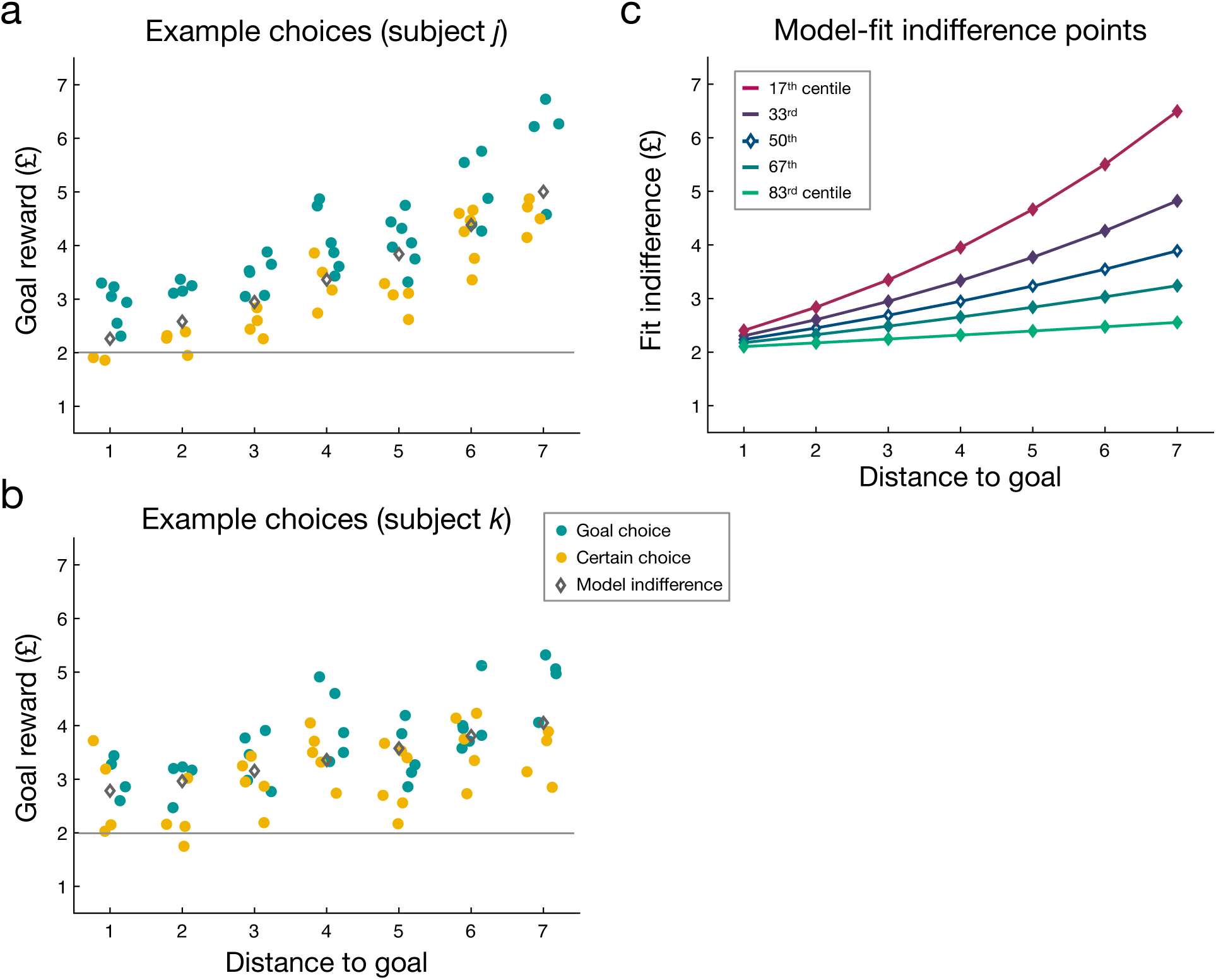
Risky goal choices and computational model fits. a-b) Choice data from two example participants with model-estimated indifference points (grey diamonds). Each dot represents a single trial; teal = goal chosen; yellow = certain option chosen. As goal distance increases, higher rewards are required to justify goal pursuit, reflecting compounding risk. Participants varied in both risk tolerance and decision noise: participant *j* (a) was more risk-averse with less variable choices than participant *k* (b). See **Figure S3** for a schematic of compounding risk effects, c) Distribution of model-estimated indifference points across distance, stratified by centiles of model-estimated step failure probability. White diamonds indicate group mean. For visualization, the intercept (goal engagement cost), was fixed at the group median.

Participants were also risk-seeking overall. When the risk-neutral expected value of the goal approximately matched the certain alternative, participants chose the risky goal 64.86 ± 1.35% of the time (t_(403)_ = 10.985, p < 0.001 versus 50%). This pattern is consistent with prior work showing that risk attitudes depend on both how uncertainty is communicated and the probability range of the risky outcome (Hertwig and Erev, 2009; Kahneman and Tversky, 1979). In particular, decisions based on brief learned experience tend to be risk-seeking than decisions based on explicitly described probabilities, especially at the probability ranges used here (Garcia et al., 2021). Our task extends this phenomenon to a setting involving extended learning across multiple sessions.

To quantify individual differences in goal pursuit, we fit a computational model in which goal values were discounted by the cumulative (exponential) risk associated with distance. The key parameter captures the subjective probability of successfully completing a single step (that is, the complement of the step failure rate). We refer to this parameter as risk-taking for goals, because higher values reflect greater willingness to pursue distant goals despite their lower expected value (**Figure S8**). The model provided a better fit than several alternatives (**Table S1**), and its step success parameter was strongly correlated with an independent, model-free estimate of risk-taking based on distance-specific indifference points (r(360) = 0.868; **Supp. Results**).

The mean step success parameter was 90.68 ± 5.98% (SD), close to but slightly higher than the experienced step success rate on day two (89.29 ± 1.91% (SD); difference t_(403)_ = 4.57, p < 0.001), again indicating mild risk-seeking.

We hypothesized that ‘Apathy–Anhedonia’ scores would be associated with reduced risk-taking for goals, paralleling its association with pessimistic subjective success ratings. However, contrary to this prediction, ‘Apathy–Anhedonia’ scores were unrelated to risk-taking for goals (t_(403)_ = −0.553, p = 0.581; **Figure 4b**), and equivalence testing ruled out even a small effect (equivalence bounds d = ±0.15; **Supp. Results**). Neither ‘Worry’ nor ‘Impulsivity’ was associated with risk-taking for goals (p-values > 0.36). A second model parameter, goal engagement cost, captured general bias toward or away from goal choices independent of distance. Exploratory analyses again found no significant relationship between this parameter and any symptom factor (p-values > 0.29), and equivalence testing again ruled out a small effect for ‘Apathy–Anhedonia’ (**Supp. Results**).

Although individuals with elevated ‘Apathy–Anhedonia’ scores reported more pessimistic beliefs about goal success, they did not show reduced willingness to pursue risky goals. At the same time, subjective success ratings and model-derived risk-taking behavior were positively correlated across participants (r(388) = 0.196, p < 0.001), indicating that beliefs and behavior were related overall even though their relationships with ‘Apathy–Anhedonia’ diverged. This raised a critical question: if that motivational symptoms were associated with pessimistic expectations but not decreased goal-pursuit, were goal-directed decisions themselves affected?

### Enhanced goal-directed behavior in apathy–anhedonia

Beyond overall risk-taking, our paradigm provided several complementary measures of goal-directed control. Across these measures, a consistent and unexpected pattern emerged: higher ‘Apathy–Anhedonia’ scores were associated with enhanced goal-directed performance, including greater sensitivity to goal value, more flexible adjustment to changes in task structure, and faster goal-directed navigation.

Our computational model included a softmax inverse temperature parameter indexing how strongly goal values influenced choice. Higher ‘Apathy–Anhedonia’ scores were positively associated with this parameter (t_(380)_ = 3.0023, p = 0.0023; **Figure 6**; **Figure S11**), indicating sharper discrimination between goal values and the certain alternative. The same pattern was observed in a model-agnostic choice quality metric, which was strongly correlated with inverse temperature (r(382) = 0.863, p < 0.001) and likewise increased with ‘Apathy–Anhedonia’ factor scores (t_(380)_ = 3.0072, p = 0.0028). ‘Impulsivity’ scores were negatively associated with inverse temperature (t_(380)_ = −2.533, p = 0.0117; **Figure 6**), although this relationship was no longer significant after controlling for location memory performance (p-values > 0.08). No significant effects were observed for ‘Worry’ (p > 0.521).

**Figure 6.**
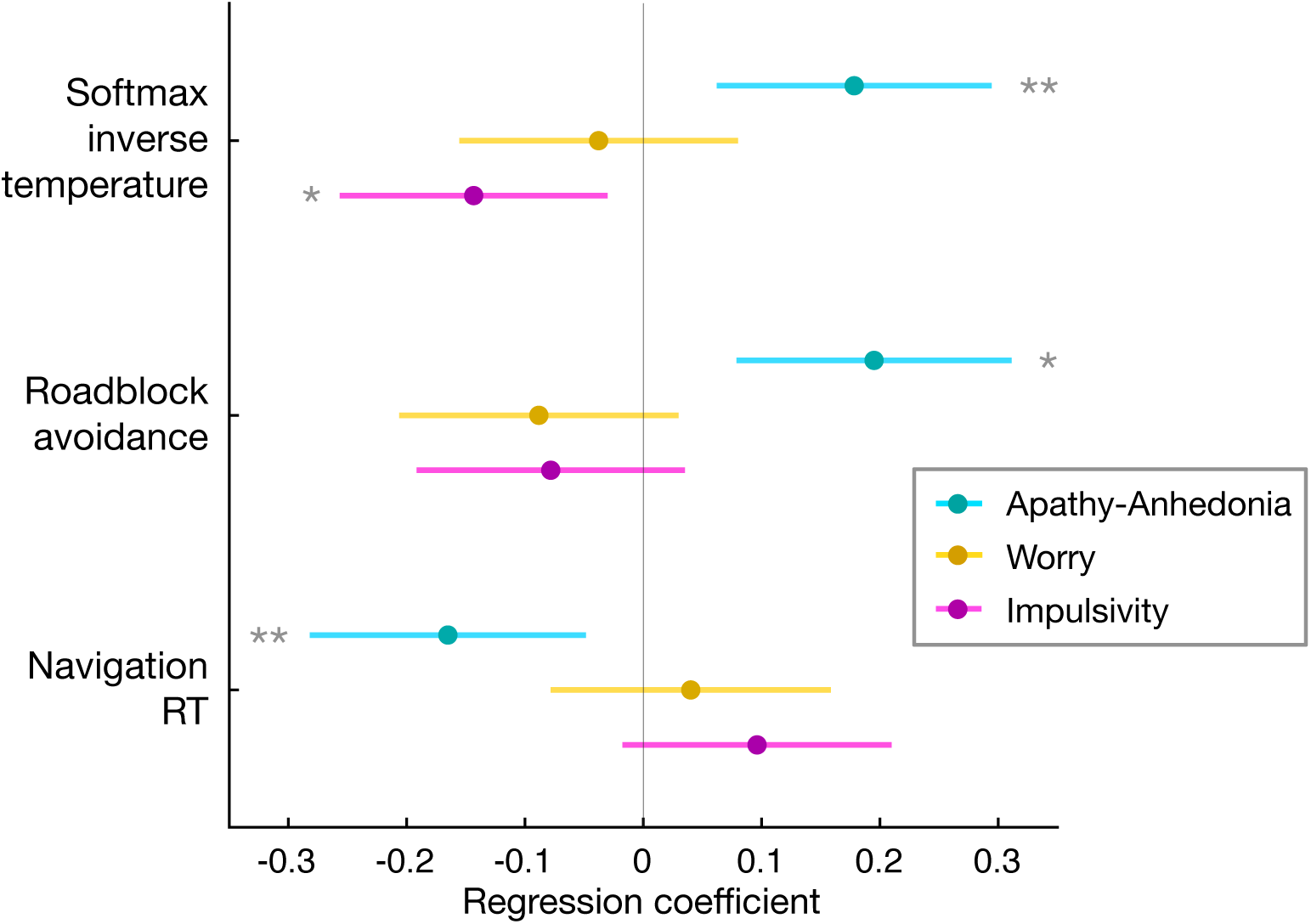
Apathy–Anhedonia is associated with enhanced goal-directed behavior. Relationship between factor scores and three measures of goal-directed performance: softmax inverse temperature (indexing the influence of goal value on choice), roadblock avoidance (indexing model-based flexibility), and navigation reaction time (indexing speed of goal-directed action). All analyses include the three transdiagnostic factors as predictors. * p < 0.05; ** p < 0.01. The association between ‘Impulsivity’ and softmax inverse temperature was not significant after controlling for location memory performance.

Occasional roadblock trials provided an independent test of model-based flexibility by probing whether participants updated their choices when a route to a goal was temporarily blocked (Tolman, 1948)(**Figure S2**). Higher ‘Apathy–Anhedonia’ scores were associated with stronger avoidance of goals when a roadblock affected the shorter path (t_(379)_ = 3.355, p = 0.0009; **Figure 6**), indicating greater model-based influence on choice. Neither ‘Impulsivity’ (t_(379)_ = −1.381, p = 0.168) nor ‘Worry’ (p > 0. 135) was significantly associated with roadblock avoidance.

A similar pattern emerged during goal navigation. After choosing to pursue a goal, participants navigated toward it via sequential keypresses. Higher ‘Apathy–Anhedonia’ scores were associated with faster overall navigation reaction times (t_(380)_ = −2.831, p = 0.005; **Figure 6**). This effect was driven by the first navigation step (t_(380)_ = −3.192, p = 0.0030, corrected), which typically determined the remainder of the path and therefore likely reflects advance planning or rapid decision-making. Reaction times on subsequent steps were unrelated to ‘Apathy–Anhedonia’ scores (t_(380)_ = −1.236, p = 0.434, corrected). Importantly, ‘Apathy–Anhedonia’ scores were not associated with first step accuracy (p > 0.91), indicating faster responses without loss of accuracy. Again, no significant relationships were observed for ‘Impulsivity’ (t_(380)_ = −1.689, p = 0.092) or ‘Worry’ (p > 0. 498).

These effects were robust across multiple control analyses. The associations between ‘Apathy–Anhedonia’ scores and value sensitivity, model-based flexibility, and navigation speed remained significant after controlling for memory performance, baseline mood, emotional responsivity to feedback, and simple trial-to-trial tendencies such as perseveration and win-stay / lose-switch behavior. Consistent with the broader pattern, higher ‘Apathy–Anhedonia’ factor scores were associated with greater reward earnings during the choice phase (t_(380)_ = 3.844, p = 0.0001; **Supp. Results**). Further, higher ‘Apathy–Anhedonia’ scores were associated with greater avoidance of goals linked to specific prior failures across a small subset of repeated start-goal locations (**Supp. Results**). Finally, convergent evidence was already present during the more limited day one session: ‘Apathy–Anhedonia’ scores predicted higher choice quality (t_(379)_ = 1.983, p = 0.0481; **Figure S12**) and faster navigation (t_(379)_ = −3.645, p = 0.003; **Figure S12**), while the roadblock effect was in the same positive direction but did not reach significance (t_(378)_ = 1.592, p = 0.112; **Figure S12**).

Given the convergence across multiple measures, we tested whether the observed relationships between ‘Apathy–Anhedonia’ and task behavior reflected a coherent multivariate pattern. We conducted a canonical correlation analysis with the three transdiagnostic factors on one side and key behavioral and belief measures on the other. The first canonical correlation was significant (r = 0.350, permutation p = 0.0002). On the symptom side, this dimension was driven by ‘Apathy–Anhedonia’ (loading = 0.803; **Figure S13**), with minimal contributions from ‘Worry’ (−0.104) or ‘Impulsivity’ (−0.138). On the behavioral side, the canonical variate captured a pattern in which pessimistic beliefs co-occurred with enhanced performance: subjective success ratings loaded negatively (−0.385), while measures of goal-directed competence loaded positively, including value sensitivity (inverse temperature), roadblock avoidance, reward earnings, and faster goal navigation responses all loading above 0.428 (**Figure S13**). In contrast, other measures loaded less than 0.137, including location memory measures, which loaded less than 0.129, confirming that the pattern was not driven by differences in environmental learning. These results demonstrate that the belief–behavior divergence associated with ‘Apathy–Anhedonia’ reflects a coherent dimension of individual variation in which pessimistic expectations coexist with sharpened value-based planning.

Taken together, these convergent findings reveal that individuals with higher ‘Apathy–Anhedonia’ scores exhibit more precise, flexible, and efficient goal-directed control, even though they report lower expectations of goal success.

## Discussion

In this study, we examined how motivational symptoms influence expectations and goal-directed decision-making in environments that require multi-step planning under learned, compounding risk. Individuals with higher transdiagnostic apathy–anhedonia scores reported lower expectations of success, yet did not reduce goal pursuit. Instead, they exhibited enhanced goal-directed behavior across multiple measures of value sensitivity, flexible planning, and navigation efficiency. Canonical correlation analysis further indicated that pessimistic beliefs and the enhanced performance measures mapped onto a shared multivariate latent factor. Together, these findings challenge the straightforward expectation that pessimistic beliefs should reduce goal pursuit, demonstrating that apathy–anhedonia was associated with more negative explicit expectations while goal-directed behavior was not merely preserved but enhanced.

The association between apathy–anhedonia and lower subjective success expectations is consistent with evidence linking depression-related symptoms to pessimistic cognitive bias (Korn et al., 2014; Mukherjee et al., 2020; Strunk and Adler, 2009). Prior work has documented pessimistic biases when uncertainty is inferred from hypothetical scenarios (Mukherjee et al., 2020; Strunk and Adler, 2009). Our results extend this pattern by showing that pessimistic beliefs also emerge when uncertainty is learned gradually through direct experience. The pessimistic bias was also stable across three days, consistent with an enduring cognitive tendency rather than a transient mood effect.

Notably, this effect was specific to motivational symptoms and was not observed for the worry factor, suggesting a dimension-specific rather than a broader distress-related bias.

Despite these pessimistic beliefs, higher apathy–anhedonia scores had no effect on risky goal pursuit. Thus, apathy–anhedonia appears to shift explicit beliefs without producing a corresponding shift in choice behavior, suggesting that symptom dimensions can decouple belief and action even within the same task context. Beliefs and risk-taking were positively correlated at the group level, confirming that the dissociation observed in individuals with elevated apathy–anhedonia cannot be attributed to the two measures tapping independent constructs. The absence of a relationship between motivational symptoms and goal pursuit is broadly consistent with evidence that depression is not reliably linked to altered risk-taking in tasks with explicitly described outcomes (Lu et al., 2024; Rutledge et al., 2017). However, this relationship has not been investigated under conditions of learned risk. This raises the possibility that pessimistic self-reports may not reflect functional impairment, a point we return to below.

The most unexpected finding was that higher apathy–anhedonia predicted not merely intact but enhanced goal-directed performance. Individuals with elevated motivational symptoms showed sharper value sensitivity in choice behavior (higher softmax inverse temperature), improved avoidance of blocked paths, more efficient navigation, and higher reward earnings. Further, convergent evidence in supplementary drift-diffusion modeling indicated that in high apathy-anhedonia, value more strongly modulated the rate of evidence accumulation (see **Supp. Results**). These effects were robust to controls for memory performance, mood, and outcome-related affect, and were unrelated to the worry factor or a measure of rumination (**Supp. Results**). Apathy-anhedonia symptoms were not significantly related to behavior in separate, simpler decision-making tasks that do not require multi-step planning (**Table S4**), including an effort-reward tradeoff task (**Supp. Results**) (Moutoussis et al., 2021).

These findings raise two main questions: first, why elevated apathy-anhedonia did not reduce goal pursuit despite more pessimistic beliefs about success; and second, why apathy–anhedonia was associated with enhanced goal-directed performance.

The finding that apathy–anhedonia was not associated with reduced goal pursuit may reflect the structured nature of the task. In daily life, apathy–anhedonia is characterized by difficulties in self-initiated goal pursuit, including generating goals, initiating action, and sustaining effort over time (Dickson and MacLeod, 2004; Le Heron et al., 2019; Scholl et al., 2022). Our paradigm bypassed many of these demands: goals were externally specified, options were explicit, and effort costs were minimal (Treadway et al., 2012). This scaffolding may neutralize the initiation and effort-allocation deficits that contribute to real-world motivational impairment, explaining why goal pursuit was not reduced. At the same time, it does not on its own explain the observed enhancement. The contrast between intact task performance and diminished real-world motivation highlights how external structure can reveal preserved competence, and may have implications for interventions such as behavioral activation that scaffold goal pursuit in clinical settings (Cuijpers et al., 2023; David et al., 2018).

At the same time, preserved goal pursuit leaves open the question of why explicit success ratings were nonetheless more pessimistic. One possibility is that ratings and choices recruited partially distinct evaluative processes. Success ratings required an absolute estimate of the likelihood of goal success in isolation, a format that places greater weight on explicit, self-referential evaluation and may therefore be especially vulnerable to pessimistic bias. In individuals with elevated apathy–anhedonia, negative expectancy and reduced reward anticipation could shift these explicit estimates downward (Hallford et al., 2020; Husain and Roiser, 2018; Korn et al., 2014). By contrast, goal choices required a bounded comparison against a concrete safe alternative, a format in which value differences may be easier to evaluate than when prospects are judged in isolation (Hsee et al., 1999). On this view, apathy–anhedonia introduces a pessimistic offset in explicit reports while leaving the comparative decision process that guides choice largely intact. This distinction may be clinically relevant, as many assessments and therapeutic contexts elicit absolute judgments of future success, which may be precisely the context in which pessimistic biases are most strongly expressed.

The most challenging finding to explain is why apathy–anhedonia was associated with enhanced goal-directed behavior. A central feature of this enhancement was sharper value sensitivity during choice, such that differences in goal value more strongly guided decisions. One possible account is that apathy–anhedonia weakens transient affective reactions during planning, shifting the balance toward more systematic integration of reward and uncertainty. Multi-step decisions require prospective simulation of possible trajectories, and those simulations are shaped not only by learned contingencies but also by the affective responses elicited by imagined outcomes (Castegnetti et al., 2020; Dolan and Dayan, 2013). If the affective impact of specific simulated outcomes is attenuated, choices may rely more directly on integrating reward magnitude with learned, compounding risk, yielding sharper value sensitivity and more precise model-based adjustment when contingencies change. This interpretation is compatible with the view that reduced reward-driven affect can promote more systematic processing (Schwarz and Clore, 2003), while locating the relevant mechanism in prospective valuation during planning.

This account is also broadly consistent with the supplementary drift diffusion analysis, in which apathy–anhedonia was specifically associated with a stronger influence of goal value on drift rate (representing evidence accumulation; **Supp. Results**). However, the mechanism remains speculative, as we have no measure of affective weighting during planning. Importantly, such a mechanism would not imply a general cognitive advantage for higher apathy-anhedonia symptoms. On this view, attenuated affective influence during planning may be beneficial when goals are externally specified, but costly in everyday settings where individuals must generate their own goals, sustain effort across delay, and tolerate uncertain feedback.

The relationship between psychiatric symptoms and goal-directed control has been most extensively studied using the two-step task. Several large-scale transdiagnostic studies have linked a dimension of compulsivity and intrusive thoughts (CIT) to reduced model-based control, while finding no such relationship for an anxious-depression dimension (Donegan et al., 2023; Gillan et al., 2016; Patzelt et al., 2019). Our impulsivity factor, which closely corresponded to CIT, was associated with weaker memory for task structure but not with goal-directed behavior after controlling for memory. A reconstructed CIT factor similarly yielded only weak or null associations with goal-directed performance (**Supp. Results**). This pattern is consistent with proposals that compulsivity-related deficits in model-based planning reflect impaired learning of task structure rather than impaired planning per se (Seow et al., 2021; Sharp et al., 2023; Sookud et al., 2025). Our multi-day design is a particular strength in this regard, because it dissociates learning from planning by assessing goal-directed behavior within a well-learned environment.

The picture for depression-related symptoms is less consistent. Smaller studies, including in clinical populations, have found some evidence of impaired goal-directed control (Gillan et al., 2020; Heller et al., 2018; Heo et al., 2021), whereas large-scale transdiagnostic studies have reported null effects (Donegan et al., 2023; Gillan et al., 2016; Patzelt et al., 2019). More recently, Sookud et al. (2025) reported enhanced model-based control at higher levels of depression symptoms. Our results go further in two ways. First, by isolating apathy–anhedonia from broader depression or anxiety dimensions, we show that the positive association is specific to motivational symptoms. Second, the association is not confined to a single model-based measure, but was evident across convergent indices including greater value sensitivity, more flexible adjustment to changing contingencies, faster goal-directed navigation, and higher reward earnings. Together, these contrasts suggest that conventional paradigms may miss symptom-specific effects on planning that become clear only in richer task environments (Scholl and Klein-Flügge, 2018; Wise et al., 2024).

These findings should be interpreted in light of several constraints. The sample was transdiagnostic rather than clinically diagnosed, although 26% met questionnaire criteria for depression and no participants were excluded based on clinical status. This dimensional approach captures symptom variation across a broad continuum and reflects real-world comorbidity. However, replication in clinical populations would strengthen confidence in these findings, including testing generalizability in patient groups with elevated apathy or anhedonia due to neurological conditions. In addition, the task used monetary incentives of moderate magnitude and involved relatively limited physical effort expenditure compared to dedicated effort-discounting paradigms (Treadway et al., 2012). Apathy–anhedonia may affect goal pursuit differently when effort costs are more salient (Bustamante et al., 2024; Treadway et al., 2012), stakes are higher, or goal pursuit unfolds over longer timescales.

In summary, motivational symptoms do not uniformly impair goal-directed behavior. In structured environments requiring multi-step planning under learned uncertainty, individuals with elevated apathy–anhedonia show more pessimistic beliefs about goal success yet enhanced goal-directed performance. These findings challenge generalized deficit models of motivation and suggest instead that apathy–anhedonia may selectively alter how expectations are represented and explicitly reported, while leaving the computations supporting value-based planning intact or, in this setting, even sharpened. More broadly, the results highlight that pessimistic self-reports about future success may not reliably index planning competence in structured tasks. If replicated in clinical samples, this dissociation suggests that environments providing external structure for goal pursuit could leverage preserved planning capacity even when explicit expectations are pessimistic.

## Methods

Participants were recruited from the online platform Prolific. Eligibility criteria included residing in the United Kingdom, being between 18–38 years old, and being fluent in English. No mental health-related screening was used. Informed consent was obtained in a manner approved by the University College London Research Ethics Committee (Approval ID No. 3451/001). Participants accessed the experiment via a web browser on desktop or laptop computers (phones and tablets were excluded). Participants were compensated at a rate of £8.21 per hour, plus performance-contingent bonus payments. The study was preregistered prior to the collection of data (https://osf.io/x3z5f).

A total of 486 participants completed both days of the risky goal task and the self-report scales. After the application of task and self-report scale exclusion criteria, 384 participants (52.2% female; mean age 29.9 years, range 18–38) with complete data remained. Participants were screened based on task performance and survey responses to (in)frequency catch questions (Zorowitz et al., 2023); see **Supplemental Methods** for details.

### Experimental task

The experiment was programmed in JavaScript using the jsPsych library (de Leeuw, 2015). The risky goal task was completed across two sessions on consecutive days. On day one, an initial instruction and practice phase familiarized participants with the task via a simple linear maze in which they explored, pursued goals, and learned about the risk of a step failure. Participants were instructed to learn item locations to support successful navigation to goals.

After practice, participants were briefly shown overhead maps of the two mazes they would later navigate (**Figure 1c**); note that these maps were shown only during breaks between blocks and not during individual trials. Participants then engaged in the exploration phase, which intermixed sets of free exploration and goal pursuit trials.

Exploration trials ended after 16 moves. Goal pursuit trials ended when the goal was reached but reward outcomes were not delivered in this phase. To measure initial learning, participants then completed an item location replacement memory test for half of the items in each maze (**Figure S1**). Next, participants completed a phase combining risk learning and goal pursuit. In this phase, they were informed that step failure would now be introduced, meaning that at each step there was a small chance of failure, which would end the trial. We did not state the specific failure probability, so participants learned the failure probability through experience. On each trial, participants were presented with a start item and a goal item (**Figure 1b**). A trial ended if the goal was successfully reached, if a step failure occurred (with an intended rate of 10%), or, rarely, if a movement response time limit expired.

To measure participants’ goal valuation and distance-related discounting, they then completed a choice phase. On each trial, participants chose between a goal option and a certain option of £2.00 (**Figure 1a**). The goal option comprised a start item, a goal item, and a goal reward value. If they chose the goal, navigation began. If they chose the certain option, the reward was displayed, followed by a delay that approximated the expected duration of goal pursuit. Goals distance varied from 1 to 7 steps (step distances were not explicitly displayed), and goal reward values increased with distance, with some random jitter around prespecified means for each distance. To estimate individual indifference points – the goal values at which participants switched between choosing the goal and the certain option – goal values for each distance were individually titrated during the choice phase (**Supp. Methods**). The choice phase also included occasional ‘roadblock’ trials. On these trials, a ‘roadblock’ object was displayed to the left of the start item with the word ‘roadblock’ below in red. This indicated that the path through that object was blocked. Some roadblocks were on the short path to the goal (‘relevant’) and some were on the long path (‘irrelevant’).

After the day one choice phase, participants completed learning phase 2, a brief set of new start-goal pairs with no certain alternative. These pairs were presented again as choice trials during day two to measure the effect of previous success or failure episodes on later decisions. Aside from these repeated trials, all other start-goal pairs were unique across the full experiment.

At the end of each session, participants completed several post-test ratings and memory measures. The first phase presented start and goal combinations. Participants rated their subjective likelihood of successfully reaching the goal on a 0-100% scale, taking into account the chance of step failure. The second measure assessed first-move accuracy, asking participants to enter the first movement direction that would lead to the goal via the shortest path. Participants then completed an item location replacement memory test, similar to that administered earlier, but with a different half of the maze locations. Finally, participants completed one generic success rating trial for each maze, in which colored squares replaced the specific start and goal items, with text indicating the goal was one step away.

On day two, the experiment proceeded as in day one, but with shorter exploration and learning phases, a longer choice phase, and no second learning phase. A location memory test was administered at the start of the session, prior to any re-exposure to the task.

### Additional tasks

Across the two main study days and a follow-up session one or two days later, participants completed additional tasks. For details, see **Supplementary Methods**. At the end of day one, participants completed a working memory measure (symmetry span). At the end of day two, a described risk task was followed by the first half of the self-report questionnaires. Participants then completed a delay discounting task, followed by the second half of the self-report questionnaires. Participants were invited to a follow-up session where they completed a test of location memory and a goal success rating phase for the risky goal task. Next, participants completed a reward learning task followed by an effort-reward decision-making task.

### Self-report measures

We collected a set of mental health questionnaires based on the set used in Gillan et al. (2016). To better target depression-related symptoms and to minimize participant fatigue, we selectively reduced or omitted less relevant scales based on findings from Hopkins et al. (2022; see also Wise and Dolan, 2020), resulting in a total of 191 items. To conservatively ensure a sufficient sample for the computational factor analysis, additional survey data were collected from a separate group of 568 participants, yielding a total of 1,028 (**Supp. Methods**).

The following scales were administered, listed in the order shown in **Figure 2**: Apathy Evaluation Scale (AES), Positive Valence Systems Scale (PVSS), Mini Mood and Anxiety Symptoms Questionnaire – Anhedonic Depression (MASQ-AD), Zung Depression Scale (SDS), Depression Anxiety Stress Scale-21 (DASS-21, referred to in **Figure 2** as ‘Internalizing’), State-Trait Anxiety Inventory (STAI), Penn State Worry Questionnaire (PSWQ), Intolerance of Uncertainty Scale – Short (IUS), State-Trait Inventory of Cognitive and Somatic Anxiety – somatic anxiety (STICSA-S), Liebowitz Social Anxiety Scale (LSAS), Barratt Impulsivity Scale (BIS), Obsessive Compulsive Inventory – Revised (OCIR), Hypomanic Personality Scale – Short (HPS). See also **Table S2**. Of these, the PVSS, MASQ-AD, DASS-21, PSWQ, IUS, STICSA-S, and HPS were added to the set of scales used by Gillan et al. (2016). Additionally, eight other items were added from other validated measures. See **Supplementary Methods** for references and details, **Tables S3–S4** for questionnaire descriptive statistics, **Figure S4** for covariance across summed scores, and **Figure S5** for histograms of scores.

### Analysis

Analyses were conducted in MATLAB 2024b, R (4.4.2), julia (1.11.5), and python. Please see **Supp. Methods** for full analysis details. Primary results were derived from the following analyses: Pearson correlations, Spearman rank correlations, linear regressions, and paired t-tests. A computational model of risky goal pursuit was developed and fit using hierarchical model fitting. Alternative models of hyperbolic discounting and models without goal engagement cost were compared using summed AIC across participants. Factor analysis was conducted using maximum likelihood estimation and oblimin rotation. Factor number was selected based on the CNG test and parallel analysis.

## Data availability

Complete behavioral data will be publicly available upon publication on the Open Science Framework.

## Acknowledgments

This research was funded in whole, or in part, by the Medical Research Council [MR/V032429/1]. For the purpose of Open Access, the author has applied a CC BY public copyright license to any Author Accepted Manuscript version arising from this submission.

## Author contributions

S.L., G.E.W., E.R, and R.R. designed the experiment. S.L. and G.E.W. collected and analyzed the data. S.L. and G.E.W wrote the analysis code. S.L., E.R., R.R. and G.E.W. contributed to data interpretation. S.L. and G.E.W. wrote the paper with E.R. and R.R.

