## Supplement for "Apathy–anhedonia is associated with pessimistic beliefs yet sharpened goal-directed planning"

#### Supplementary Methods

#### Supplementary Results

1. Transdiagnostic factor analysis
2. Learning, memory, and impulsivity
3. Risk-taking for goals: null relationship to factors
4. Risk-taking for goals and risk-taking propensity
5. Risk-taking for goals and model-free measures
6. Goal-directed behavior and alternative factors
7. Goal-directed task earnings and Apathy-Anhedonia
8. Goal-directed behavior: null relationship with rumination
9. Reward modulation of drift rate in HDDM and Apathy-Anhedonia
10. Risk-taking for goals and additional tasks
11. Additional tasks and transdiagnostic factors
12. Mood at baseline
13. Emotional responses to outcomes
14. Additional preregistered hypotheses

#### Supplementary Figures & Tables

### Supplementary Methods

#### Participants

Our target sample size after applying exclusion criteria was 400 participants, providing greater than 99% power to detect a 'small' effect size of Cohen's d = 0.2 (α = .05, two-tailed, approximate Pearson r = 0.10). A total of 510 participants began the first session. Of these, 486 participants completed both sessions of the risky goal task and the self-report questionnaires, which were administered on day two. Following the application of task-related and survey-related exclusion criteria, 384 participants remained (52.2% female; mean age 29.9 years, range 18-38). Exclusion criteria led to the removal of 82 participants based on task behavior and 26 participants based on survey catch question performance, leading to a combined exclusion of 102 (21% of the sample; criteria were non-exclusive). See subsequent section for exclusion criteria details. The main experiment consisted of two sessions conducted on consecutive days. Participants were invited to complete a follow-up session, to be completed on day three or four. For this session, 372 participants with complete behavioral data and self-report data were included in analyses.

Primary analyses relating factors to behavior used the sample of 384 participants meeting task and survey criteria. Some behavior-only analyses used a larger set of 404 participants with complete task data but incomplete or excluded survey data. For the day 3 follow-up session, the corresponding number was 385.

For the transdiagnostic factor analysis of the self-report scales, data from 460 participants with complete survey data were included; these additional participants either had missing risky goal task data or did not pass risky goal task exclusion criteria. To increase the sample size, 568 additional participants were recruited, resulting in a total sample of 1,028 for factor analysis. The total sample size was determined based on a conservative guideline of a five-to-one ratio of participants to items in factor analysis (Gorsuch, 1983). The factor analysis results were stable when the sample size was halved. The additional participants completed different tasks in addition to the scales, and as with the primary sample, they were recruited for a two-day study. This was a deliberate choice, as pilot data in other studies indicated that participant exclusion rates due to poor performance were higher when recruiting for only a single-session study.

Participants were not screened for additional health or mental health conditions. The Prolific platform collected such information only through optional additional questions, which many potential UK participants declined to answer in 2023-2024. Specifically, in separate online studies in this period, we found that adding any voluntary health or mental health screening questions led to an approximately 50% reduction in the available participant pool. The remaining participants who opted to complete the additional health questions exhibited significantly lower depression and anxiety scores compared to participants recruited without additional screening. Thus, to include a larger group of participants with higher depression and anxiety scores, we did not include additional screening questions. Finally, we did not collect data about mental health diagnoses or treatments in the study, as this level of personal data was not permitted by the study's ethics approval.

#### Experimental task details

The task consisted of sequential phases across two consecutive days. See **Figure 1d** for a visual timeline, in which the width of each arrow indicates the relative duration of each phase. The primary focus of the preregistered analyses was the choice phase on day two. All start-goal combinations were unique across the task, with the exception of 10 specific combinations that were intentionally repeated (described below); repeated combinations were excluded from all primary analyses. Participants were assigned to one of three counterbalancing orders, each corresponding to a different prespecified mapping of object images to maze locations.

Day one included the following phases in this order: practice, exploration, location memory test, learning, choice, second learning, and post-tests. Day two included the following phases in this order: location memory test, exploration, learning, choice, and post-tests.

Although the overall structure was similar across days, the choice phase on day one was half the length of that on day two. Analyses focus on the day two choice phase, as this followed extensive learning and overnight consolidation. Additionally, on day two, the location memory test was given immediately at the start of the session, and the subsequent exploration and learning phases were shorter.

Participants navigated the mazes using the keyboard arrow keys. At each location, only two directions led to new locations; these valid directions were visually indicated by the absence of a “wall” on the corresponding edge. Pressing an arrow key toward a blocked edge (i.e., hitting a “wall”) resulted in no movement. Participants were free to choose their route, including retracing their steps or turning around. If a response (choice or directional input) was not made within 15 seconds, the trial ended with a warning. The learning and choice phases included risk, in which every move – including those that hit a wall – carried the same fixed probability of triggering a step failure event, which terminated the trial. The per-step failure probability was fixed and did not vary with participant behavior. During navigation, the identity of the current maze could be inferred from the start and goal objects as well as a small image of the maze’s distinctive rock background in the upper-right corner of the screen (**Figure S1a**, right). This maze identity indicator was not shown on the decision screen on choice phase trials.

On day one, participants began with a combined instruction and practice phase using a simplified non-looping maze. This phase comprised three parts: free exploration, goal-directed navigation, and the introduction of step failure risk. Participants were instructed that the best strategy was to learn object locations in order to plan the shortest path to goals. The instructions included a general example to illustrate how more steps increased the probability of step failure. Participants were also told that the same mazes would be presented again on day two, to emphasize the value of learning and remembering the mazes. Before proceeding, they completed a brief comprehension quiz; correct answers were required to advance to the main task.

Following the instructions and practice, we collected self-report measures of fatigue and time awake. For the fatigue item, participants responded to the question “How tired are you?” by moving a circle along a slider bar with the labels “Very energetic”, “Moderately energetic”, “Neither tired nor energetic”, “Moderately tired”,

“Extremely tired”. For the hours awake item, participants were asked “How many hours have you been awake today?” and responded with numeric values.

To accelerate learning in the main task, participants were presented with an overhead view of the two mazes they would be navigating. One maze was taller and the other was wider, and each was displayed against a distinct rock background image (**Figure 1**). Participants could switch between the mazes using the arrow keys. The minimum total viewing time across both mazes was 10 seconds. These overhead maps were presented again briefly during breaks between main task phases, but they were not shown during individual trials of the task.

Before each main phase, participants responded to the question “How happy are you right now?” by moving a circle along a slider bar anchored at “Very unhappy” and “Very happy”.

**Exploration phase.** In the exploration phase, participants learned the maze structure through direct experience. There were 12 trials in total on day one, divided into two types: pure exploration trials and goal-directed trials with no risk of failure. In pure exploration trials, participants were shown an object representing their starting location, with the text “Explore to learn” displayed to the right. After 16 exploration steps (with wall hits also counting as moves), “Good work” appeared in blue text beneath the current location object.

At the start of each trial, participants selected an initial direction. If a response was not made within the 15-second time limit, a warning (“Please respond faster! −0.50 GBP”) was displayed in red text, and the trial ended. After selecting a direction, a confirmation arrow replaced the object image during a 0.25 s inter-stimulus interval (ISI), after which the next location object appeared. In the goal-directed trials, participants were presented with a start location object and a goal location object to the right (**Figure 1**), with the word “goal” presented below (in place of the reward amount used in later phases). In these trials, the ISI between object presentations was 1.25 s, and this same ISI was used for all goal-directed navigation trials. These trials ended after reaching the goal, which could take as many steps as necessary. After reaching the goal, “Good work” was presented beneath the current location object.

Exploration trials were ordered as follows: four exploration trials, two goal-directed trials, two exploration trials, and four goal-directed trials. The goals for the goal-directed trials were all 5–6 steps away from the starting location, and maze identity was balanced across these trials. After the last trial, participants were asked “Do you wish to explore more?” If they selected ‘yes’, they were given one more exploration trial in each maze. This process could repeat until participants chose to move on. Few participants selected this option more than once.

On day two, the exploration phase followed a location memory test (described below). It consisted of 10 trials: six exploration trials and four goal-directed trials. The first two exploration trials had a move limit of 16; the remaining four had a move limit of 12.

**Location memory test.** After the exploration phase on day one, participants completed an object location replacement memory test. This measure was repeated at the end of day one, at the start of day two, and at the end of day two. On each trial, participants were shown a single object picture above a schematic grid of the corresponding maze, with locations labeled a–p (**Figure S1b**). Maze identity was indicated by the shape and width of the grid; no background rock image was shown. Participants indicated the object’s location by pressing the corresponding letter key. After a response, the screen elements were replaced with a confidence scale, on which participants rated their response confidence from low (1) to high (4). There was no time limit for memory or confidence responses. The 32 maze locations across the two mazes were divided into two sets, with each set tested once per day. Objects were shown in a pseudo-random order.

**Learning phase.** To allow participants to learn the risk associated with goal pursuit through extended experience, the learning phase introduced the risk of step failure into goal-pursuit trials. (Step failure had been introduced during practice.) This phase also provided continued exposure to maze structure. Goal-pursuit trials were structured as described above in the exploration phase, with three differences: first, each goal was associated with a reward value that could be added to participants’ earnings; second, each navigation step was associated with a 10% chance of failure, independent of participants' actions; third, participants were required to press the "space" key three times after entering their direction response. The learning phase comprised 32 trials on day one and 12 on day two.

In each trial, participants navigated toward the goal, whose image remained on screen (**Figure 1**). Participants entered the direction of navigation using the arrow keys, after which they pressed the "space" key three times to continue. The space-key response had a time limit of 7.5 seconds; exceeding this limit terminated the trial. This was followed by a 1.25 s ISI during which the object disappeared and was replaced by an arrow representing the selected direction. If navigation to the goal was successful, "Collected: £ [amount]!" was shown below the goal image in orange text for 2.0 s. A jittered ITI of 2.25 s (1.5–3 s range) followed each trial. In the event of a step failure, the central image was replaced with an image representing falling, "Step fail!" was shown below in red text, the goal image was removed, and the missed reward value was changed to grey text for 2.0 s (**Figure 1**). To balance the duration of successful and failed goal-pursuit trials, after failure, participants waited for the amount of time that would have been required to finish navigation to the goal (approximated as the number of remaining steps to the goal multiplied by 2.5 s, based on pilot data).

**Choice phase.** After gaining experience with goal-directed navigation and step failure risk, participants started the choice phase. Here, participants chose between goal options of varying value and a certain alternative, allowing estimation of individual risk preference. The choice phase comprised 30 trials on day one and 70 on day two. The goal option consisted of a start location, goal location, and goal reward value. The certain alternative was always £2.00. The goal and certain options were presented in the upper and lower halves of the screen (**Figure 1a**; **Figure S1a**), with screen position randomized across trials. Participants could indicate their choice using the up and down arrow keys, with a 15 s time limit. If a response was not recorded, a warning "Please respond faster! -0.50 GBP" appeared in red text, and the trial ended. If the goal option was selected, the display transitioned to the navigation screen (**Figure 1b**; **Figure S1a**, right) displaying the start, goal, and goal reward value. The trial then proceeded as in a learning phase trial. If the certain option was selected, the display was replaced with “Collected certain: £ 2.00!” in orange text for 2 s. To match the expected duration of goal pursuit, this was followed by a delay proportional to the number of steps along the shortest path to the goal.

The choice phase included occasional ‘roadblock’ trials to assess flexible goal-directed decision-making. On these trials, an additional image was displayed to the left of the start item with the word ‘roadblock’ below in red. This indicated that the path through that location was blocked. On half of the roadblock trials, the roadblock was placed on the relevant shorter path to the goal. On the other half, the roadblock was on the irrelevant longer path to the goal (and thus could be ignored). Participants were not told which path the roadblock affected; this had to be inferred from knowledge of the maze structure. A roadblock on the shorter path effectively transformed a normal goal located 4 steps away into a goal located 12 steps away (shifting the expected success probability from 66% to 28%). The shortest-path distance on roadblock trials was fixed at 4 steps (not disclosed to participants). To encourage engagement with the roadblock trial offers, the base goal reward value was multiplied by a factor of 1.14–1.17. The choice phase on each day contained two relevant and two irrelevant roadblock trials. We constructed an index of roadblock avoidance as the difference in goal choice rate between irrelevant and relevant roadblock trials.

For further details about the composition of the choice phase trial list, online adjustment of goal offer values, adjustment of step failure probability, and post-outcome emotion ratings, see the section **Choice phase implementation** below.

**Second learning phase.** On day one, following the choice phase, participants completed a non-choice learning phase. This phase included specific start-goal combinations that would be repeated on day two to assess the influence of prior success or failure episodes on choice. (No other start-goal combinations were repeated in the experiment.) For trial structure, see the learning phase section above. This phase included 12 trials. The first and last trials were ‘buffer’ trials, which were not included on day two. In the 10 trials of interest, the success or failure of goal pursuit was predetermined, unlike all other trials. Participants were not informed of this. In these trials, the natural stochastic step failure process was disabled. For six trials, a step failure event was programmed to occur at a specific step location (e.g., step two). For the remaining four trials, participants reached the goal successfully. The rate of step failure across all steps in this phase remained close to the target rate of 10%. On day two, these 10 trials were pseudo-randomly interspersed among the standard choice trials. To better capture any influence of the prior episode, on day two, the goal reward value was set to be near the base value for a given distance (factor of 0.98–1.02).

**Success probability rating and memory tests.** At the end of the main session on each day, we collected measures of learning and memory. The first phase assessed learning about step failure risk by collecting subjective success probability ratings for specific start-goal combinations. Participants were shown a display similar to the learning phase trial format, showing a start and goal location but no goal value, with a rating scale below (**Figure S1c**). On each trial, participants were asked "What is your expected probability of goal success?" The rating scale ranged from 0–100%, with labels at 0% ("Never"), 50% ("Half"), and 100% ("Always"). In the instructions for this phase, participants were reminded to consider the possibility of step failure events. This phase included 14 trials in a pseudo-random order, with one trial per distance per maze. No time limit was placed on responses. A 1.0 s ITI followed each trial. The start-goal combinations in this phase were novel.

The second phase measured memory for the direction of the first move along the shortest path to the goal, given a novel start-goal combination. Participants were shown a display similar to the learning phase trial format, showing a start and goal location but no goal value, with arrow icons and the text "Press first key to best reach goal" below. This phase included eight trials, with one trial per maze for goal distances of 3–6 steps. No time limit was placed on responses. A jittered ITI (mean 1.75 s, range 1.0–2.5 s) followed each trial.

The third phase measured location memory, as described above, testing 16 locations that were not tested at the beginning of the day.

The fourth phase was an exploratory measure that assessed subjective estimates of success probability for generic rather than specific start-goal combinations.

On each trial, two blue squares replaced the start and goal location item images. The start image contained the text “any Start”, while the goal image contained the text “Goal 1 step from start” above one left-facing arrow. This phase included two trials, assessing success ratings for a one-step goal in each of the two mazes. On day two, this phase was expanded to include additional distances (3 and 6 steps for one maze; 4 and 7 steps in the other), in addition to the distance-1 trials.

**Post-session questionnaire.** At the end of each session, participants completed a brief questionnaire assessing several aspects of their experience. Items included self-ratings of task engagement, ability to minimize distractions, and understanding of key task components. Participants also reported their confidence in remembering object locations, described their decision-making strategies, and provided any additional comments. On the second day, participants were also asked whether they used external aids (such as images of the mazes) and to rate their overall task enjoyment. Finally, participants answered questions about their current stress level and recent sleep quantity and quality.

**Memory and subjective success ratings – day three follow-up.** On the morning of the third day, participants received an invitation to participate in an additional session. This session was also announced at the end of day two. This session, which could be completed the same day or the following day, included a location memory test and a subjective goal success rating phase. Each phase followed the same procedures as described above. For the subjective goal success rating phase, new start-goal combinations were used for each maze and distance.

**Choice phase implementation**. In this section, we describe the choice phase trial list composition, the adjustment of goal offer values and step failure probability, and the inclusion of post-outcome emotion ratings.

Across learning and choice phases, the trial lists were approximately balanced across key factors, including equal representation of the two mazes, different starting locations in each maze, different goal locations in each maze, clockwise or counterclockwise directions of navigation along the shortest path, and the direction of the first move in navigation.

The base goal values across all trials were determined by pilot data and adjusted throughout the task, as described below. For distances 1–7, the values (in GBP) were: 2.76, 2.94, 3.16, 3.40, 3.68, 4.02, and 4.41. These values were close to the risk-neutral expected value (reward magnitude multiplied by the cumulative step success probability) plus a small goal engagement cost representing the time and effort of navigation. For example, the expected value at distance 7 was £4.18, while the list value was £4.41.

To estimate individual differences in risk-taking for goals (indifference points), presented goal values were adjusted in two ways. First, for normal choice trials (excluding roadblock trials and repeated trials from day one), goal values were jittered using a fractional multiplier. The multiplier was drawn approximately uniformly from the range [0.734, 1.266]. These values were pseudo-randomly assigned to trials to maintain balance across distances and mazes.

Second, the goal values were titrated online to approximate each participant's distance-specific indifference points. This was implemented via a participant- and distance-specific scale parameter, which was multiplied by the base goal value to produce the offer displayed on each trial. At the start of the day one choice phase, the parameter was initialized to 1.0 at each distance. The scale parameter was updated following each choice in which the jitter multiplier was moderate (0.8–1.2; see above). The scale parameter was decreased following goal choices and increased following certain choices, with an update increment that started at 0.09 and decayed as a function of trial number. For example, if the certain option was chosen on the first three distance-5 trials, the scale parameter would reach approximately 1.27, increasing the goal value presented on the next distance-5 trial. To accelerate convergence, the scale parameter for a given distance was also updated following choices at neighboring distances, with the neighbor updates set to one-third of the primary increment for distances 2–6 and two-thirds for the extreme distances (1 and 7), where fewer neighboring trial types were available. Simulations confirmed that this procedure successfully converged to distance-specific indifference points, such that offers were distributed around each indifference point with variation determined by the jitter multiplier. Critically, the procedure did not assume a specific functional form for discounting across distance, allowing individual differences in the shape of distance-based discounting to be captured. Scale parameter values from the end of day one were carried over to the start of day two.

To minimize the impact of stochastic variation on the experienced step failure rate, the task dynamically adjusted the per-step failure probability when the observed rate diverged from the 10% target. Step failures and total steps were tracked separately for each maze across the learning and choice phases. When the step failure rate exceeded an upper tolerance bound, the probability was temporarily reduced to 5%; when it fell below a lower bound, the probability was temporarily increased to 30%. The tolerance window narrowed across the task: ±2.75% for trials 20–40, ±2.225% for trials 40–60, and ±2.0% for trials beyond 60. On day two, an adjustment was applied on 28.2% of trials on average across participants.

To measure emotional responses to goal outcomes and to test potential symptom associations, occasional post-outcome happiness ratings were collected during the learning and choice phases (Rutledge et al., 2014). Ratings used the same graded happiness scale described above for between-phase measurement. Several ratings were collected during the learning phase to provide initial experience. During the choice phase, the task aimed to collect five ratings following goal successes, five following step failures, and three following certain-outcome choices. Ratings were triggered after every three success events, three failure events, or four certain choices. If the target number of ratings had not been reached within 20 trials (e.g., due to infrequent goal choices or an imbalanced success/failure rate), the sampling frequency for the underrepresented outcome type was increased. Two exploratory frustration ratings were collected, using the prompt “How **frustrated** are you right now?” with anchors “Not at all frustrated” and “Very frustrated.” One was triggered after the fourth goal success and the other was triggered after the fourth step failure.

#### Participant exclusion and data quality

As noted above, participants were excluded based on task behavior and survey performance. Task performance was screened using seven criteria (described below), leading to the removal of 82 participants from behavioral analyses (16.9% of the 486 who completed both sessions). Five of seven task-based exclusion criteria were informed by pilot data and prespecified in the preregistration (accounting for 76 of 82 exclusions). Applying the two additional exclusion criteria (accounting for six participants) did not qualitatively alter any of the reported results. Further preregistered criteria were also specified, but no participants met them. Of the 82 excluded for task behavior, eight also failed self-report scale exclusion criteria. The criteria were not mutually exclusive. The criteria and the number of participants excluded for each criterion were as follows:

1. Poor memory: Low accuracy on the maze object location replacement memory test, defined as less than 66.7% correct on the test at the end of day two (chance = 50%). Exclusion of 27 participants (5.6%), 10 of whom were also excluded for at least one other criterion.
2. Poor first-direction accuracy: Poor memory for direction combined with poor location memory. Specifically, participants were excluded if they met all three of the following conditions: less than 66.7% accuracy on first-direction moves during the choice phase (chance = 50%), less than 66.7% accuracy on first-direction moves during the end test phase and less than 75% accuracy on location memory at the end of day two. Exclusion of four participants (0.8%), one of whom was also excluded for at least one other criterion.
3. Strong choice bias: Choosing either the goal option or certain option on 85% or more of trials overall or in either half of the day two choice phase. Exclusion of 35 participants (7.2%), 16 of whom were also excluded for at least one other criterion.
4. Outlier navigation behavior: Extremely high rates of wall hits during navigation, defined as more than 100 wall hits within a given day. Note that walls are clearly marked by black boundaries. This criterion was relaxed from the preregistration threshold of > 2 SD above the mean wall hit rate. In the data, this primarily appeared as more than 100 wall hits during the day one exploration phase, whereas most other participants had fewer than five. Exclusion of eight

participants (1.6%), three of whom were also excluded for at least one other criterion.

5. Poor value sensitivity during choice: The average value of chosen goal options was less than or equal to the value of rejected goal options. A ‘choice quality’ measure, used in later analyses, was computed by first z-scoring the offered goal values within each distance. We then subtracted the mean goal value for non-chosen goals from the mean value of chosen goals and averaged these values across distances to yield a single measure for each participant. (Note that this measure empirically approximates a logistic regression coefficient but is more stable with low trial numbers.) With zero as the cutoff value, the mean in the retained sample was 1.080 ± 0.363 (range 0.031 – 1.63). Exclusion of 20 participants (4.1%), 14 of whom were also excluded for at least one other criterion.
6. Missed choices: Missing more than 20% of trials in the day two choice phase (indicating low attention). Exclusion of seven participants (1.4%), five of whom were also excluded for at least one other criterion.
7. Strong aversion to short-distance goals: Strong avoidance of short-distance goals relative to medium-distance goals, combined with choice quality below 0.5. This pattern reflects a U-shaped relationship between goal value acceptance rate and distance. Relative avoidance was defined using the end-stage goal value scaling parameter during the task (see below), where the scale at distance 1 was 0.3 higher than the scale at distance 4. Exclusion of three participants (0.6%).

Among the included participants, a small number of trials were excluded based on excessively fast response times. For choices between goal and certain options on days one and two, trials were excluded if the reaction time was less than 400 ms, a threshold determined from the data distribution. The aim was to retain as much data as possible while removing trials too fast for adequate processing of choice information. This resulted in the exclusion of a mean of 0.024 trials per participant (range 0-6) on day one and 0.026 trials (range 0-6) on day two.

Next, participants were excluded from symptom-related analyses based on poor performance on (in)frequency (catch) items embedded in self-report scales, as recommended by Zorowitz et al. (2023). This led to the exclusion of 26 participants (5.3%). Of these 26 excluded for survey performance, eight also failed task-based exclusion criteria described above.

We defined exclusion criteria based on errors in responses to the nine catch items. Of these, four items were treated as strict catch items: “I breathe every day”; “I know the months of the year”; “I have been to the moon”; “I know how to count to ten”. The remaining five items were evaluated less strictly (e.g., “I am sometimes suddenly completely physically invisible to cameras”), as higher rates of errors indicated that some participants may have interpreted these items differently due to ambiguous wording. Errors were converted into percentage deviation from the expected correct endpoint (e.g., “Always” or “Never”). The exclusion criteria were as follows: 1) answering two or more strict catch items by any deviation from the correct response, 2) missing two or more catch items overall by more than 34% (generally more than 1 scale step away from the correct response), or 3) answering three or more catch items overall by any deviation from the correct response.

#### Self-report measures

To examine relationships between goal-directed behavior under uncertainty and depression symptoms, we used a transdiagnostic approach to capture factors that cut across traditional diagnostic categories. Previous transdiagnostic approaches have commonly identified a combined anxious-depression factor rather than separate factors (e.g., Gillan et al., 2016). To better distinguish depression from anxiety symptoms, we included several additional scales (see below) in addition to those used by Gillan et al. (2016). To minimize participant burden, less critical scales were shortened based on item reduction results from Hopkins et al. (2022; see also Wise and Dolan, 2020), which identified the minimal items sets needed to reproduce the compulsivity and intrusive thought (CIT) factor. The schizotypy and alcohol use scales were omitted because Hopkins et al. (2022) found that these scales were not critical for replicating the factor structure, and the symptoms assessed were not of preregistered interest. Two eating disorder scale items that contributed to the reduced CIT factor in Hopkins et al. (2022) were also omitted. The administered set of scales included 191 items. Scales assessing real-world risk-taking propensity were also included for comparison with task-based risk preferences.

To minimize fatigue, the scales were divided into two halves. Scale order was randomized for each participant. The self-report battery was administered after the risky goal task on day two. Before each half, participants were instructed that the scales were as important as the preceding risky goal task and were asked to maintain their attention throughout. The first half was administered after a separate described-risk task. Participants then completed a delay discounting task, which served as a break, before the second half of the scales.

The following mental health questionnaires were administered (**Table S2**): Apathy Evaluation Scale (AES) (Marin et al., 1991), Positive Valence Systems Scale (PVSS) (Khazanov et al., 2020), Mini Mood and Anxiety Symptoms Questionnaire – anhedonic depression (MASQ-AD) (Clark and Watson, 1995), Zung Depression Scale (SDS) (Zung, 1965), Depression Anxiety Stress Scale-21 (DASS-21) (Lovibond and Lovibond, 1995), State-Trait Anxiety Inventory (STAI) (Spielberger et al., 1983), Penn State Worry Questionnaire (PSWQ) (Kertz et al., 2014), Intolerance of Uncertainty Scale – Short (IUS) (Carleton et al., 2007), State-Trait Inventory of Cognitive and Somatic Anxiety – somatic anxiety (STICSA-S) (Ree et al., 2008), Liebowitz Social Anxiety Scale (LSAS) (Liebowitz, 1987), Barratt Impulsivity Scale (BIS) (Patton et al., 1995), Obsessive Compulsive Inventory – Revised (OCIR) (Foa et al., 2002), and Hypomanic Personality Scale – Short (HPS) (International Personality Item Pool). Of these, the PVSS, MASQ-AD, DASS-21, PSWQ, IUS, STICSA-S, and HPS were added to the set of scales used by Gillan et al. (2016).

To ensure data quality, we employed (in)frequency (‘catch’) questions, as recommended by Zorowitz et al. (2023), who demonstrated that traditional instructed catch questions were less successful at detecting low-quality data in online studies. For example, one infrequency item was: "I have been to the moon." Nine (in)frequency catch questions were included, with one item in each of the following scales: AES,

PVSS, SDS, DASS-21, STAI, PSWQ, IUS, BIS, and a risk-propensity scale (DOSPERT).

**Additional mental health items.** Additional items from other depression and anxiety-related scales were included if the item content was not represented by an existing item in the above scales and if that content seemed either central to clinical definitions of depression or anxiety or relevant for the current study. For the Patient Health Questionnaire-9 (PHQ-9) and Generalized Anxiety Disorder-7 (GAD-7) in particular, one additional item from each was added as the content was not included in the other scales. In total, eight additional items were added from seven different scales. The items were added to the scale whose instructions and response format most closely matched the source scale.

Patient Health Questionnaire-9 (PHQ-9) (Kroenke et al., 2001): item 5 was appended to the end of the DASS scale: "I have a poor appetite or overeat."

Generalized Anxiety Disorder-7 (GAD-7) (Spitzer et al., 2006): item 7 was appended to the end of the STAI scale: "I feel afraid, as if something awful might happen."

Hospital Anxiety and Depression Scale (HADS) (Zigmond and Snaith, 1983): items 2 and 6 were appended to the end of the MASQ-AD scale: "I feel as if I am slowed down" and "I have lost interest in my appearance."

Fatigue Severity Scale (FSS) (Krupp et al., 1989): item 9 was appended to the end of the AES scale: "Fatigue interferes with my work, family, or social life."

Beck Depression Inventory (BDI) (Beck et al., 1961): item 5 was appended to the end of the STAI scale: "I do not feel particularly guilty."

Revised Life Orientation Test (LOT-R) (Scheier et al., 1994): item 1 was appended to the end of the STAI scale: "In uncertain times, I usually expect the best."

Rumination-Reflection Questionnaire (RRQ) (Trapnell and Campbell, 1999): item 3 was appended to the end of the STAI scale: "I tend to 'ruminate' or dwell over things that happen to me for a really long time afterward."

**Selection of items from scales.** For the primary depression and anxiety scales (AES, PVSS, MASQ-AD, SDS, DASS-21, STAI), all items from these scales or subscales were retained, as were the short forms of the HPS and IUS.

The MASQ anhedonic depression subscale was taken from the Mini-MASQ (Clark and Watson, 1995). The subscale is composed of eight items, of which six are positively valenced (see also use by Gagne et al., 2020). The items included, based on numbering from the Mini-MASQ, were: 1, 5, 9, 11, 15, 19, 23, 25. This subscale overlaps substantially with the MASQ-D30-AD (Wardenaar et al., 2010), a more recent 10-item subscale composed only of positively valenced items. Five of the six positive Mini-MASQ-AD items overlap with MASQ-D30-AD items, and studies have found high correlations between these measures (Corral-Frías et al., 2019; Lin et al., 2014).

For the SDS, to improve response accuracy, the first response option was modified from “A little of the time” to “Rarely or none of the time”, based on this usage in the German and Dutch SDS translations. This modification was intended to better accommodate participants with low depression scores, particularly for the suicidality item. In support of this modification, an alternative adaptation of the SDS (Guy, 1976) similarly added a “None of the time” option.

The PSWQ was shortened based on an 8-item short form (Hopko et al., 2003) and the analysis of Kertz et al. (2014). Item 12 (“always worrying”) was removed due to redundancy with item 7 (“all my life”), given the instructions to rate how typical or characteristic each item was for respondents. Item 7 was retained based on its higher loading onto a single factor (Kertz et al., 2014). Item 14 was included to capture uncontrollable worry, aligning with its inclusion in a validated 3-item short form (Berle et al., 2011). The final 8-item selection (numbered from the full scale) included: 2, 4, 5, 6, 7, 9, 13, 14.

For the STICSA somatic anxiety scale, four items overlapped with DASS items (STICSA items: 1, 8, 15, 18; DASS: 2, 4, 7, 19). These items were omitted from the STICSA but were compensated for in scoring. For scoring, the four overlapping DASS item scores were substituted for the omitted STICSA items when computing the STICSA total.

The LSAS was reduced following Hopkins et al. (2022) as this scale was aligned with our primary interest in core depression and anxiety symptoms. Given the unitary structure of the scale, the item set was reduced further based on factor loading strength reported by Oakman et al. (2003). The final 13-item selection (numbered from the full scale) included: 2, 5, 7, 8, 10, 11, 12, 14, 16, 18, 19, 20, 23. Only the fear/anxiety ratings were collected, omitting the avoidance ratings, to minimize response burden.

The BIS was reduced following Hopkins et al. (2022) and prior psychometric studies that evaluated eight-item reduced versions (Morean et al., 2014; Steinberg et al., 2013). The two eight-item versions comprised identical items. We supplemented these items with additional CIT factor items identified by Hopkins et al. (2022). The final 13-item selection (numbered from the full scale) included: 1, 2, 5, 6, 8, 9, 12, 13, 14, 17, 19, 22, 25.

The OCIR was reduced following Hopkins et al. (2022) to include only CIT factor items. The final 10-item selection (numbered from the full scale) included: 2, 3, 4, 6, 7, 9, 12, 13, 15, 18. After completion of data collection, inspection of the supplementary item weights from Hopkins et al. (2022) revealed a discrepancy between the supplemental item weights and the item list in the main tables, such that OCIR items 11 and 16 also had non-zero weights. These items were not included in our questionnaire set; however, in Hopkins et al. (2022), item 11 received the second-lowest CIT factor weighting and item 16 received an intermediate weighting.

**Risk-taking propensity measures.** We measured general risk-taking via the General Risk Propensity Scale (GRIPS) (Zhang et al., 2019) and the general item from the General Risk Question (GRQ) (Dohmen et al., 2011; Frey et al., 2017). The GRIPS is an 8-item scale in which each item assesses attitudes toward risk or perceptions of risk in general. The General Risk Question (GRQ) (Dohmen et al., 2011) was first used in the German Socio-Economic Panel (SOEP). We adopted a 6-item version, with a general item and one item each for driving, financial matters, leisure and sports, work or study, and health. We also included items from the Domain-Specific Risk-Taking Scale (DOSPERT; Weber et al., 2002), specifically those items related to ethics, health-safety, and recreation.

#### Additional tasks in order of collection

**Working memory measure – symmetry span (day one).** After completing all risky goal task phases on day one, participants completed a task targeting working memory (WM). We adapted a commonly used operations span (OSPAN) task, replacing arithmetic operations with visual symmetry judgments (SymSPAN) (Kane et al., 2004), given that variability in arithmetic abilities may add unwanted variance. As in the original OSPAN, participants were required to maintain a series of presented letters in memory. This differed from the original symmetry span, which used spatial location memory (Kane et al., 2004). Participants were shown an 8 x 8 black-and-white grid and judged whether the left and right halves were symmetrical. After a response, a to-be-encoded letter appeared (e.g., B), followed by the next symmetry judgment. Letter sequences ranged in length from three to eight. At the end of a sequence, participants were asked to type the letters in order presented, with no time limit. The task consisted of eight trials. Sequence lengths of four, six, seven, and eight were each used once, while lengths three and five were each used twice. Trial order was determined so that difficulty increased across the task but not monotonically. Each trial used unique symmetry images and letters in pseudo-random order. To ensure adequate practice before the main task, participants received detailed instructions followed by staged practice on each sub-task individually and then combined. Scores were calculated by summing the number of letters in sequences that were recalled entirely correctly, divided by the total number of letters presented (Wimmer et al., 2018). The mean letter score was 62.8 ± 21.2% SD (range 0–100%). Participants were excluded from WM analyses if their performance on symmetry judgments was two standard deviations below the mean. As expected, the mean symmetry performance was high (98.2 ± 6.9% SD).

**General intelligence (day one).** Following the working memory task, participants completed six items from the International Cognitive Ability Resource (ICAR; Condon and Revelle, 2014), following Dubois and Hauser (2022), as a coarse measure of individual differences in general intelligence. In practice, however, several features of the testing context compromised the validity of this measure. The ICAR was administered at the end of the longest session in the study, when participants were likely to be fatigued, consistent with open-ended feedback. In addition, unlike all other tasks in the study (besides working memory), the ICAR offered no performance-contingent monetary incentives and lacked a repetitive, task-engaging structure, both of which may have reduced sustained effort. Under these conditions, variation in ICAR performance was likely to reflect nuisance factors such as persistence, motivation, and conscientiousness rather than general cognitive ability itself. For this reason, we did not include ICAR scores in any analyses.

**Described risk prospects (day two).** After completing all phases of the risky goal task on day two, participants completed a described risk prospects task. We adopted a commonly used risk task (Rutledge et al., 2014), assessing risk preference in described one-shot decisions in two conditions: (1) a risky option with gain or zero outcomes versus a certain alternative; (2) a risky option with mixed gain and loss outcomes versus a certain alternative. (A loss-only condition was not included due to time constraints and because losses were less related to the main risky goal task.) Participants completed 24 trials in each condition, where trials from both conditions were pseudo-randomly intermixed across the task. On each trial, participants chose between a risky option that offered equal probabilities of a better or worse outcome versus a certain option. Accumulated points contributed to participants' monetary bonus. Participants first completed two practice trials: one from the mixed condition and one from the gain condition. The risky option was always on the left, the certain option on the right. Participants used the 'g' key to select the risky option and the 'h' key to select the certain option. If participants chose the risky option, there was a 1.3 s delay and then the gamble outcome was displayed for 1 s. If they chose the certain option, the outcome was shown for 2 s. Trials were followed by a 2 s ITI. In the mixed condition, the risky option involved a potential gain (ranging from 50 to 80 points across 3 gain levels) or loss (–90 to –13 points across 8 loss levels), whereas the certain option always yielded 0 points. In the gain condition, the certain option provided a fixed reward (30 to 50 across 3 levels), while the risky option offered a 50% chance of receiving a larger reward (60 to 158 across 8 levels) and a 50% chance of receiving 0 points. The first half of the self-report scales was administered after this task.

**Delay discounting (day two).** After the first half of the survey scales on day two, participants completed a delay discounting task. In this task, participants chose between a smaller amount of money available that day ("today") versus a larger amount of money to be received at a delay of 1–180 days. The immediate option values ranged from £1 to £42, while the larger later amount was either £25 or £45. There were 30 trials in total. Participants were instructed that one choice would be randomly selected to count for real, and that a fraction of this amount would be added to their bonus payment. The offers were created based on the approach of Lempert et al. (2020). The offers were designed to estimate hyperbolic discounting rates ranging from k = 0.00043 to k = 0.2525, equally spaced in log(k) space. Additional offers targeted k values of 0.0105 and 0.0171 (medium discounting), while the k = 0.00071 level (very low discounting) was omitted. Immediate reward values and delays were combined to achieve approximately uniform coverage of the reward-by-delay space, after scaling for the two delayed amounts. The first trial was a practice trial. Options were presented on the left and right side of the screen, with screen position randomized across trials, and participants entered their choice by using the left and right arrow keys. The trial order was pseudo-randomized. To simplify bonus payments for a large sample, the bonus was always based on a randomly selected trial where the immediate option was chosen. The second half of the self-report scales was administered after this task.

**Reward learning (day three follow-up).** On day three, following the risky goal task follow-up measures, participants completed a reward learning task. The task was a two-armed bandit with fixed stimulus-reward associations. Participants learned to select the more frequently rewarded stimulus in each pair (78% and 22% reward probability). Each of six stimulus pairs was presented 18 times in pseudo-random order, yielding 108 trials. Stimulus-reward assignments were counterbalanced across three lists, one of which was randomly assigned to each participant. Performance contributed to a bonus payment. Before the main task, participants completed six practice trials. On each trial, two choice options were presented to the left and right of the screen center, with left/right locations randomized. Participants selected the left option by pressing the ‘f’ key and the right option by pressing the ‘j’ key. The response window was 3 s. After the response, the chosen option was highlighted for 1.0 s, followed by a blank inter-stimulus interval of 3.0 s. Feedback for the chosen option was then presented for 1.5 s, followed by a jittered ITI (mean 2.25 s, range 1.5–3.0 s) with a fixation cross.

**Effort–reward decision-making (day three follow-up).** After the reward learning task on day three, participants completed an effort-reward tradeoff task. We adopted a widely used paradigm for examining trade-offs between monetary reward and physical effort: the Effort-Expenditure for Rewards Task (Berwian et al., 2020; Treadway et al., 2012). On each trial, participants chose between a “hard task” that required more key presses (40–130) to inflate a virtual balloon and yielded more points (4–7), and an “easy task” that required fewer key presses (20–70) and yielded fewer points (3–6). The effort and reward levels were varied over 62 trials in pseudo-random order, with the harder option always offering more points than the easier option. Participants used the ‘d’ key to select the left option and the ‘f’ key to select the right option. Due to a coding error, the response window was 2 s rather than the intended 6 s; however, if no response was recorded, the same trial was repeated. This error may limit the interpretability and generalizability of any effects observed in this task. Participants then inflated the balloon by pressing the ‘j’ key, with a maximum duration of 40 s to complete the required key presses. If presses were not completed, the trial was repeated. A 2 s ITI followed each trial. On 22 trials, selected at random and without advance notice, participants were not required to execute the effort component of the trial after making their choice. Performance contributed to a bonus payment.

#### Analysis

Data processing was conducted with custom scripts written in Matlab (2024b), and general analyses were conducted using the Matlab functions corr for bivariate analyses (Pearson or, when the distributions were highly skewed, Spearman), regstats for basic regression analyses, and partialcorr for Spearman partial correlations. Where applicable, null effects were tested using the two one-sided tests (TOST) procedure implemented with the tsum_TOST function in the TOSTER package in R. Equivalence bounds, specified in terms of effect size, were set to rule out a "small" effect size or greater (Cohen's d = 0.20; Pearson r = 0.100), below which effects were considered equivalent to zero. The included sample size provided 97.5% power for the equivalence test.

#### Factor analysis

We performed exploratory factor analysis (EFA) on the self-report questionnaire data. 191 items went into the factor analysis. As specified in the preregistration, our aim in constructing the questionnaire battery was to differentiate depression from anxiety. Following the procedures of Gillan et al. (2016) and Hopkins et al. (2022), we used the factanal() function in R, selecting maximum likelihood estimation (MLE) and oblique rotation (oblimin) to allow correlation between factors. We selected the number of factors based on Cattell's criterion (Cattell, 1966) and used the Cattell-Nelson-Gorsuch (CNG) test to objectively identify the 'elbow' in the scree plot using the nCng() function (Cattell et al., 1981). We also used Parallel Analysis following Hopkins et al. (2022), and both the CNG test and Parallel Analysis supported a three-factor structure. Seven items were removed because they did not load at ± 0.25 or above on any factor, leaving a total of 184 items (Gillan et al., 2016).

#### Behavioral analysis

Memory location replacement accuracy was computed as a scaled metric from 0 to 1, where 0.5 represents chance. Accuracy was calculated as (maximum error – mean distance error) / maximum error. In the 16-location loop mazes used here, the maximum possible error was eight steps and the expected error at chance was four steps.

To examine basic influences on decision-making, we tested the effect of goal value on choice in a multilevel logistic regression model, with random intercepts for distance and subject, using the glmmTMB function from the glmmTMB package in R. To examine the effect of distance, participant- and distance-specific indifference points were estimated using logistic regression. The effect of distance on indifference point was then analyzed in a multilevel regression model, with distance and subject as random effects, using the lme function from the nlme package in R.

In the risky goal task choice phase, to characterize individual risk-seeking or risk-averse behavior, we used a model-free strategy that examined behavior on decisions where a risk-neutral discounted goal offer was close to the value of the certain offer (a probability-discounted goal value between £1.90–2.10). A risk-neutral agent would be expected to show no preference. A bias toward choosing the goal option on these trials would indicate risk-seeking, and vice versa.

We computed a 'choice quality' measure based on the influence of goal offer value on decisions. Unlike the computational models of decision-making (below), this metric makes no assumptions about the form or presence of discounting and allows for arbitrary shifts in indifference points across distances. At each distance, the goal offer values for all trials were z-scored. The mean z-scored goal value for the rejected goals was then subtracted from the mean for the chosen goals. These values were then averaged across the seven distances to create a single choice quality measure for each participant, where values above zero indicate a positive influence of goal offer value on choices to pursue the goal. This measure approximates a logistic regression coefficient when trial counts per distance are sufficiently large.

We computed a model-free estimate of risk-taking for goals using a simple method to estimate distance-specific indifference points. Within each participant, for each distance, if the goal was chosen on more than two trials, the median goal value for rejected offers was subtracted from the median goal value for accepted offers. This method provided similar results to an alternative based on logistic regression but avoided instability from extreme regression coefficients.

Goal success probability ratings, collected at the end of days one and two, were screened to remove participants who appeared to rate failure probability instead of success probability. This inverse pattern was relatively rare: on day two, 92% of participants exhibited the expected negative correlation between success rating and distance (median $r = -0.889$). Exclusion was based on a positive correlation between distance and rating, with tolerance for cases where the range of ratings was very restricted. Specifically, participants were excluded from this analysis if 1) the correlation between rating and distance was greater than 0.5 and 2) the probability rating for distance 1 plus a tolerance of 0.05 was less than the rating for distance 7. This resulted in the exclusion of two participants from this analysis on day one, eleven participants on day two, and two participants on day three.

The same criteria were applied to the exploratory measure of success ratings for general goals at a given distance (with no specific start or goal location indicated). The procedure was adjusted to use the ratings for distances 6 and 7, because only one maze was rated at each distance. Additionally, on day one, only distance 1 ratings were collected, so no exclusions were applied. These data were not collected on the day three follow-up.

#### Computational model of decision-making for risky goals

To model individual differences in risk preferences for uncertain goals, we fit two classes of computational models to participants' choices. Both models assume that participants decide by comparing the subjective value of pursuing the risky goal, $V_{risky}$, to the value of the certain offer, $V_{certain}$.

$$P(choose\ risky) = logit^{-1}(\beta(V_{risky} - V_{certain})) \quad (1)$$

Here, $\beta$ is an inverse temperature parameter (free, varying across participants) that governs choice stochasticity.

For both models, $V_{certain}$ includes both the (constant) reward of taking the certain offer (£2) and a subjective bonus amount (c) reflecting avoidance of an additional cost to navigating the maze, referred to as goal engagement cost, $V_{certain} = 2 + c$. $c$ is a free parameter allowed to vary across participants. Models without $c$ were also tested and we found that its inclusion improved model fit in all cases.

The models differ in how $V_{risky}$ is computed on each trial. In the exponential discounting model, $V_{risky}$ is the probability of successfully reaching the goal times the trial-varying reward, $m$, that would be received:

$$V_{risky} = m \times (1 - p)^d \tag{2}$$

The probability of reaching a goal at a given distance is the success probability $p$ (one minus the failure probability) raised to the power of the trial-varying distance $d$. We treat $p$ as a free parameter that varies across participants, reflecting individual differences in the subjective assessment of per-step success probability.

In the hyperbolic discounting model, $V_{risky}$ is computed by discounting the maze reward, $m$, according to a hyperbolic function:

$$V_{risky} = \frac{m}{1 + kd} \tag{3}$$

Here, $k$ is a free parameter that varies across participants specifying the extent of hyperbolic discounting with distance, $d$.

For each participant, we estimated the free parameters of each model using an expectation maximization (EM) procedure (Huys et al., 2011) that maximizes the likelihood of choices, implemented in the Julia language (Bezanson et al., 2017)(code from: https://github.com/ndawlab/em). This procedure maximizes the likelihood of each participant's choices, treating individual parameter estimates as random effects drawn from group-level Gaussian distributions whose means and covariances are jointly estimated. This approach allowed partial pooling of parameter estimates across individuals, which helped constrain extreme estimates while retaining subject-level flexibility.

We compared models using three metrics. We computed integrated Akaike Information Criterion (iAIC) and integrated Bayesian Information Criterion (iBIC) for each model. These were computed by defining the marginal likelihood of the data given either model aggregated across subjects, marginalizing per-subject parameters with the Laplace approximation, and penalizing for the group-level parameters using either AIC or BIC (Huys et al., 2012). We additionally computed unbiased per-subject marginal likelihoods via subject-level leave-one-out cross-validation (**Table S1**).

To assess the identifiability and reliability of model parameters for the best-fitting model, we conducted a parameter recovery analysis (Wilson and Collins, 2019). For each participant, we simulated synthetic choice data using the fitted parameters. We then refitted the same model to the simulated data using identical fitting procedures. Recovered parameters were compared to the generating parameters to evaluate recovery accuracy. Parameter recovery, estimated via the correlation between original parameter values and recovered values, was high: success probability ($r(382) = 0.926$); goal engagement cost ($r(382) = 0.912$); softmax inverse temperature ($r(382) = 0.894$).

#### Canonical correlation analysis

To test whether the observed associations between symptom dimensions, expectations, and goal-directed performance reflected a coherent multivariate pattern rather than independent effects, we conducted a canonical correlation analysis (CCA) using the 'canoncorr' function in Matlab. On the symptom side, we entered the transdiagnostic factor scores (Apathy–Anhedonia, Worry, Impulsivity). On the behavioral side, as variables of interest, we entered success ratings, softmax inverse temperature, roadblock avoidance, (log) goal navigation step response time, repetition of previous successful goal, avoidance of previous failed goal, and reward earned. We also entered location memory performance, estimated per-step failure probability, estimated goal engagement cost, and (log) choice reaction time. All variables were z-scored prior to analysis. Statistical significance of the canonical correlations was assessed using a permutation test (10,000 permutations) in which the mapping between symptom and behavioral matrices was randomly shuffled to generate a null distribution. We report the first canonical variate pair, which captures the linear combinations of symptom and behavioral variables that maximize the correlation between the two sets.

#### Supplemental drift diffusion model of decision-making for risky goals

We used hierarchical Bayesian drift diffusion modeling (HDDM) to estimate latent decision parameters from participants' trial-level risky choices (Wiecki et al., 2013). Models were implemented via the dockerHDDM environment (Pan et al., 2025), a standardized, containerized platform that ensures reproducibility across systems. Models were fit to participants' trial-by-trial decisions and reaction times to test whether reward influenced the drift rate. Following prior work (Ort et al., 2019), we used informative priors (Wiecki et al., 2013) and varied only the drift rate parameter (v) interactions across nested models, holding other parameters constant. We estimated a baseline model and a model including an interaction between drift rate and reward (goal value).

All models were estimated using Markov Chain Monte Carlo (MCMC) sampling with four chains of 2500 samples each (1000 discarded as burn-in). Convergence was assessed using Gelman–Rubin $\hat{R}$ statistics and visual inspection of traces. The baseline model failed to converge (max $\hat{R} = 1.17$). The model with the reward interaction converged adequately ($\hat{R} = 1.03$); thus, this was the model used.

#### Analysis of additional tasks

**Described risk prospects.** Exclusion criteria were applied to ensure data quality. First, participants were excluded if more than 25% of their choices had outlier reaction times (see criteria below; n = 2), or if their choices were too noisy, defined as choices not systematically influenced by the values of the presented options (gain: n = 8; mixed: n = 10). We calculated a choice quality measure similar to that used in the risky goal task. In the gain condition, the widely varying gain option was z-scored after adjusting for the three certain option levels, and the mean z-scored value for rejected prospects was subtracted from mean for chosen prospects. In the mixed condition, the widely varying loss option was z-scored after adjusting for the three gain option levels. As with the risky goal task, this measure approximates a logistic regression coefficient with sufficient data. Among participants with at least 10% variability in their choices, those whose choice quality fell below the group mean minus 2 standard deviations were excluded. For included participants, individual trials were excluded if reaction times were outliers, defined as follows: faster than 300 ms (based on inspection of distribution), slower than 20 seconds, or slower than the participant's mean reaction time plus 2.5 standard deviations. These same criteria were used to determine the >25% outlier-based participant exclusion.

In the computational model of the risk task, the risk-aversion parameter (α) and the loss-aversion parameter (λ) determine utility of each component, scaled by its respective probability:

$$U_{gamble} = 0.5(V_{gain})^{\alpha} - 0.5\lambda {V_{loss}}^{\alpha} \tag{4}$$

$$U_{certain} = (V_{certain})^{\alpha} \tag{5}$$

First, the two conditions were modeled together to estimate a more stable softmax inverse temperature parameter. Second, the two conditions were modeled separately to estimate risk and loss parameters for correlations with self-report risk propensity. When modeled separately, in the gain condition, the loss aversion parameter did not apply. The softmax function from Equation 1 was used to calculate the probability of choosing the risky versus certain option based on their subjective values. Parameters were estimated with maximum likelihood using Matlab's fmincon function.

**Delay discounting.** Exclusion criteria were applied to ensure data quality. First, participants were excluded if more than 25% of their choices had outlier reaction times (see criteria below; n = 3), or if their choices were too noisy, defined as choices not systematically influenced by the values of the presented options (n = 5). Participants were excluded if their behavior was not adequately captured by the computational model (see below). Based on visual inspection, this threshold was set to a summed log-likelihood value of less than -7.5 (where lower values indicate worse fit). For included participants, individual trials were excluded if reaction times were outliers, defined as follows: faster than 500 ms (based on inspection of distribution), slower than 20 seconds, or slower than the participant's mean reaction time plus 2.5 standard deviations. These same criteria were used to determine the >25% outlier-based participant exclusion described above.

We used a standard hyperbolic discounting model (e.g., Enkavi et al., 2019). Here, the value of the delayed option (A) is discounted by increasing delay (D) of reward receipt in a hyperbolic manner, where the free parameter *k* represents an individual's discount rate:

$$V = \frac{A}{1 + kD} \tag{6}$$

The softmax function from Equation 1 was used to calculate the probability of choosing the delayed versus immediate option based on their subjective values. Parameters were estimated with maximum likelihood using Matlab's fmincon function.

**Reward learning.** Exclusion criteria were applied to ensure data quality. First, participants were excluded if greater than or equal to 20% of their choices were missed (n = 0). Second, participants were excluded if more than 14% of their choices were faster than 300 ms (n = 3). Third, participants were excluded if accuracy (choice of the higher-reward option) over the last five repetitions was less than or equal to 50% correct (n = 29). Application of these non-exclusive criteria led to the exclusion of 30 participants (8%). Performance was measured as the overall mean accuracy across the task. In this task, as fixed reward associations prevented the identifiability of a softmax parameter separate from a learning rate parameter, overall mean performance was used as a 'choice quality' measure.

**Effort-based decision-making.** Exclusion criteria were applied to ensure data quality. First, participants were excluded if more than 25% of their choices were too fast (n = 0), or if their choices were too noisy, defined as choices not systematically influenced by the values of the presented options (n = 3). We calculated a choice quality measure similar to that used in the risky goal task. First, we converted the combined button press and point offers into one value per option by computing ratios of presses per point for each option. The mean z-scored value for rejected options was subtracted from the mean of chosen options. As in the risk-taking tasks, this measure approximates a logistic regression coefficient with sufficient data. Among participants with at least 10%

variability in their choices, those whose choice quality fell below the group mean minus 1 standard deviation were excluded. A stricter threshold was used in this task given the coding error that restricted the response window to 2.0 s. For included participants, individual trials were excluded if reaction times were faster than 250 ms (based on inspection of distribution).

In the computational model of the effort task (Berwian et al., 2020), the value of option *a* is determined by the trade-off between reward *r*(*a*) and effort *e*(*a*) scaled by separate sensitivity parameters:

$$V(a) = \beta_{rew} * r(a) - \beta_{eff} * e(a) \tag{7}$$

The softmax function from Equation 1 was used to calculate the probability of choosing the high-effort versus low-effort option based on their subjective values. Parameters were estimated with maximum likelihood using Matlab's fmincon function.

### Supplementary Results

#### 1. Transdiagnostic factor analysis

The first factor was characterized by positive loading on scales related to apathy and anhedonic depression (AES, M = 0.55 ± 0.17 (SD); MASQ-AD, M = 0.65 ± 0.12) and negative loading on the scale assessing positive valence symptoms (PVSS, M = -0.54 ± 0.09). The second factor was characterized by positive loading on scales related to worry (reduced PSWQ, M = 0.71 ± 0.02), intolerance of uncertainty (IUS, M = 0.65 ± 0.05), and social anxiety (reduced LSAS, M = 0.50 ± 0.04). The third factor was characterized by positive loading on the impulsivity scale (reduced BIS, M = 0.44 ± 0.14), as well as weaker positive loading from the scales related to mood instability (HPS, M = 0.39 ± 0.06) and obsessive-compulsive symptoms (reduced OCIR, M = 0.32 ± 0.09).

As hypothesized, the inclusion of additional scales effectively differentiated a depression-related factor ('Apathy-Anhedonia') from an anxiety-related factor ('Worry'). The top-loading items on the first factor included AES item "I have motivation," followed by MASQ items "I felt really lively, 'up'" and "I felt really happy." The top-loading items on the second factor came from the PSWQ, including "I know I shouldn't worry about things, but I just cannot help it" and "When I am under pressure I worry a lot." The top-loading items on the third factor came from the BIS, including "I act on impulse" and "I act on the spur of the moment."

The third factor resembled the 'compulsivity and intrusive thoughts' (CIT) factor identified previously (Gillan et al., 2016). A derived CIT factor based on the weightings from Hopkins et al. (2022) was strongly correlated with 'Impulsivity' factor scores ($r(382) = 0.826$; **Figure S4**). However, the 'Impulsivity' factor had relatively higher loading from impulsivity items and relatively lower loading from obsessive-compulsive items compared with the weights in Hopkins et al. (2022). Our battery included 24 of 26 items in the reduced CIT factor in Hopkins et al. (2022); the omitted items came from an eating disorders scale. The derived CIT factor was nearly identical to a CIT factor derived using the weightings of Wise & Dolan (2020) ($r > 0.99$).

Although the factors were derived from exploratory factor analysis, we also found that each factor could also be approximated well by a weighted combination of the three strongest contributing scales in the included participant sample. ‘Apathy-Anhedonia’ was correlated at r = 0.975 with a weighted combination of summed AES (w = 1), MASQ-AD (w = 1.5), and PVSS (w = –0.25). ‘Worry’ was correlated at r = 0.980 with a weighted combination of summed IUS (w = 1), PSWQ (w = 1.5), and LSAS (w = 0.6). ‘Impulsivity’ showed a lower but still strong correlation (r = 0.909), as expected given the broader distribution of contributing items, with a weighted combination of summed BIS (w = 1), HPS (w = 0.7), and OCIR (w = 0.375).

Risk propensity measures were not included in the factor analysis because they assess behavioral tendencies rather than mental health symptoms. General risk propensity scores were moderately correlated with factor scores, ranging from r = –0.307 (‘Worry’) to r = 0.275 (‘Impulsivity’).

#### 2. Learning, memory, and impulsivity

In this section, we first summarize learning and memory performance and then present additional results related to the ‘Impulsivity’ factor and a derived CIT factor.

In the item location memory test, performance increased across day one (early 83.9 ± 13.9% (SD); late 91.7 ± 9.7%; $t_{(399)} = 15.199$, $p < 0.001$; **Figure 3a**). On day two, location memory performance also increased across the session (early 91.2 ± 9.6%; late 95.3 ± 6.4%; $t_{(401)} = 11.417$, $p < 0.001$). There was no significant change in memory from the end of day one to the start of day two ($t_{(400)} = -1.382$, $p = 0.168$).

The majority of participants also returned for a follow-up session (day three), one to two days after the main risky goal task. Location memory performance remained high in this follow-up session (93.8 ± 8.2%). While performance was lower than at the end of day two ($t_{(384)} = -5.016$, $p < 0.001$), it was higher than at the start of day two ($t_{(384)} = 7.982$, $p < 0.001$).

First-move direction accuracy during goal navigation in the choice phase was high on day one and increased further on day two (86.5 ± 16.4%; 91.2 ± 8.7%; $t_{(397)} = 6.484$, $p < 0.001$). First-move direction accuracy varied across distances to the goal. In particular, direction accuracy was lower for goals one step away than for goals at a middistance (four steps) away (one step 88.9%; four steps 93.9%; $t_{(382)} = -5.484$, $p < 0.001$), suggesting some difficulty discriminating between adjacent locations in memory. Direction accuracy measured in separate end-of-session probe questions was consistently lower than performance during active navigation (day one mean 75.5%; day two mean 87.0%; both significantly lower than accuracy during the choice phase, p-values < 0.0001); thus, our analyses focused on behavior during the choice phase.

Based on previous research (Gillan et al., 2016; Seow et al., 2021; Sharp et al., 2023; Sookud et al., 2025), we hypothesized that a factor related to compulsivity and intrusive thoughts (CIT) would be negatively associated with the acquisition of model-based knowledge of the maze (**hypothesis 14**). Our 'Impulsivity' factor, which positively weighted items related to intrusive thoughts, most closely aligned with the prior CIT factor. As noted above, a derived CIT factor based on the weightings from Hopkins et al. (2022) was highly correlated with 'Impulsivity' factor scores across participants ($r(382) = 0.826$; **Figure S4**).

As described in the **Results** section, regression analyses revealed that the 'Impulsivity' factor was significantly negatively associated with location memory across days (**Figure 3**). These associations were confirmed in bivariate analyses of the individual memory variables (**Figure S6**).

We also found that 'Impulsivity' factor scores were negatively related to first-move direction accuracy in goal pursuit during the choice phase on both days (day one $r_s(377) = -0.106$, $p = 0.0385$; day two $r_s(380) = -0.105$, $p = 0.0411$; partial correlations). One possible interpretation of this effect is that lower direction accuracy reflected weaker goal-directed (model-based) behavior. However, the relationship with 'Impulsivity' factor scores was better accounted for by location memory performance, as including location memory in the model rendered the association with direction accuracy non-significant (day one $r_s(376) = -0.0324$, $p = 0.530$; day two $r_s(379) = -0.0499$, $p = 0.331$), while location memory remained significant (day one $r_s(376) = -0.130$, $p = 0.0116$; day two $r_s(379) = -0.107$, $p = 0.0375$). This suggests that 'Impulsivity' factor scores were more strongly related to poor location memory than to impaired goal-directed navigation.

Further analysis of direction accuracy also supported the interpretation that fine-grained location memory underlies the relationship with ‘Impulsivity’ factor scores. As noted above, direction accuracy was lower for goals one step away than for more distant goals. We found that the relationship between ‘Impulsivity’ factor scores and direction accuracy was driven by accuracy for goals directly adjacent to the start location (distance one: $r_s(379) = -0.142$, $p = 0.0054$, partial correlation). The difference in direction accuracy between distance 1 and distance 4 was also associated with ‘Impulsivity’ scores ($r_s(379) = -0.136$, $p = 0.0079$).

Finally, in exploratory analyses, we examined the relationship between the derived CIT factor and memory. Although correlations with memory were uniformly negative, CIT factor scores were not significantly related to memory performance in bivariate analyses (p-values > 0.12; **Figure S6**). When we included a derived anxious-depression (AD) factor and the LSAS scale as covariates, to approximate the approach of Gillan et al. (2016), we again found no significant effects (p-values > 0.26).

These results offer mixed support for our hypothesis that the factor most closely related to the previously identified compulsive and intrusive thought factor would relate to building a memory representation environmental structure. Our ‘Impulsivity’ factor was the factor most closely related to the previous CIT factor (**Figure S4**) and showed consistent negative associations with memory. However, a derived CIT factor based on a reduced set of items using weights from Hopkins et al. (2022) showed no significant correlations with memory.

#### 3. Risk-taking for goals: null relationship to factors

As reported in the Results section, we found no associations between ‘Apathy-Anhedonia’ factor scores and risky goal pursuit choices (model-estimated success probability). Here, we used the two one-sided tests (TOST) procedure to examine whether even small effects could be ruled out, allowing us to test whether the effect was statistically equivalent to zero. We tested against equivalence bounds of effect size $d = –0.15$ and $d = 0.15$, tighter than the conventional small effect threshold ($d = 0.20$). With 384 participants, we had 80% power to test an effect of this size with the TOST procedure. These analyses were based on bivariate regression results, because the TOSTER package in R does not support multivariate models.

We found that the relationship between ‘Apathy-Anhedonia’ factor scores and risky goal pursuit was statistically equivalent to zero (TOST equivalence test $t_{(383)}$ = -1.669, p = 0.0480; coef = 0.0648, 90% CI [-0.0194, 0.149]), indicating that we can rule out any effect stronger than d = 0.15. The effect for the ‘Worry’ factor was also equivalent to zero (‘Worry’ $t_{(383)}$ = -1.674, p = 0.0475; coef = 0.0645 CI [-0.0197, 0.1487]). The effect for the ‘Impulsivity’ factor was not equivalent to zero ($t_{(383)}$ = -1.460, p = 0.0726; coef = 0.0754 CI [-0.0085, 0.1592]), indicating that we could not rule out effects as small as d = ±0.15.

Similarly, the relationship between ‘Apathy-Anhedonia’ factor scores and the goal engagement cost parameter was statistically equivalent to zero within bounds of d = 0.15 (TOST p = 0.0019), and the relationships with the other factors were also statistically equivalent to zero within bounds of d = ±0.15 (TOST p-values < 0.04).

#### 4. Risk-taking for goals and risk-taking propensity

Separately from the transdiagnostic analysis, we hypothesized that risk-taking for goals would be positively related to self-reported risk-taking propensity measures. This was the case for one propensity measure (GRQ general; $t_{(379)}$ = 2.657, p = 0.0164, controlling for transdiagnostic factors, corrected for two comparisons) but not the other (GRIPS $t_{(379)}$ = 0.530, p > 0.99, corrected). To provide an effect size comparable with prior studies (Frey et al., 2017; Steiner and Frey, 2021), we also estimated a bivariate correlation between the GRQ general item and risk-taking for goals alone, yielding r(382) = 0.113 (p = 0.0265, uncorrected). A similar correlation was observed for the summed GRQ score (r(382) = 0.112, p = 0.0277).

For subjective ratings of goal success, self-reported general risk-taking propensity was significantly related to ratings on day one only (GRQ general $t_{(377)}$ = 2.352, p = 0.0384; GRIPS $t_{(377)}$ = 2.661, p = 0.0163; corrected for two general risk measures). On day two, this relationship was in the same positive direction but not significant (GRIPS: p = 0.052; GRQ general: p = 0.107; corrected for two comparisons).

#### 5. Risk-taking for goals and model-free measures

As a complementary model-free measure of risk-taking for goals, we examined choice indifference points at different distances using a simple median difference between accepted minus rejected goal values. Given the compounding risk, we expected that indifference points at longer distances, adjusted for general goal preference, would best approximate risk-taking preference. We therefore used the mean indifference point for distances 6 and 7 goals. To account for a general baseline tendency to pursue or avoid goals, we subtracted the mean of the indifference points at distances 1 and 2. Here, higher indifference values indicated lower risk-taking. These values could not be estimated for all participants.

The model-free risk measure was highly correlated with the model-derived risk-taking for goals ($r(342) = -0.869$, $p < 0.001$). Overall, the median group indifference point estimated across distances closely tracked the risk-neutral expected value of goals across distances, with no deviation on average (calculated using each participant's experienced per-step success rate; $t_{(403)} = -1.411$, $p = 0.159$). When we examined indifference points at shorter and longer distances separately, we found significant risk aversion at shorter distances (mean of distances 1–2; $t_{(390)} = 6.420$, $p < 0.001$, corrected for two comparisons) and risk-seeking for longer-distance goals (mean of distances 6–7; $t_{(373)} = -5.099$, $p < 0.001$, corrected for two comparisons). These analyses included only participants with estimable indifference points at the relevant distances. Turning to the model-free measure of risk-taking, replicating the pattern observed for the model-derived risk-taking measure, no symptom factors were significantly correlated with the model-free measure, including the 'Apathy-Anhedonia' factor ($p > 0.96$). Consistent with the model-derived results, the GRQ general item was significantly associated with the model-free measure, with higher indifference points for distant goals corresponding to lower self-reported risk propensity ($r(342) = -0.114$, $p = 0.035$; GRIPS $r(342) = -0.0497$, $p = 0.358$).

A second model-free measure of risk-taking for goals was available directly from the task, in the form of scaling parameters for each goal distance that were adjusted throughout the task to balance rates of goal and certain choices. This measure, derived from the online titration procedure, relies on additional assumptions relative to the indifference point measure. Similar to the indifference point measure, we took the mean scale values for distances 6 and 7 and subtracted the mean scale values for distances 1 and 2. We found that the scale parameter measure was highly correlated with model-derived risk-taking for goals ($r(382) = -0.924$, $p < 0.001$). This measure also replicated the significant correlation with the GRQ general item. Both alternative measures yielded results consistent with the model-derived risk-taking parameter.

#### 6. Goal-directed behavior and alternative factors

In this section, we first report relationships between the 'Worry' and 'Impulsivity' factors and goal-directed behavior. Second, we report relationships between derived compulsivity and intrusive thoughts (CIT) and anxious-depression (AD) factors and goal-directed behavior.

The three measures of goal-directed behavior that were positively associated with 'Apathy-Anhedonia' scores (softmax inverse temperature, roadblock avoidance, and navigation response time) were moderately intercorrelated. Softmax inverse temperature was correlated with roadblock avoidance ($r(381) = 0.355$, $p < 0.001$) and with (log) navigation response time ($r(382) = -0.337$, $p < 0.001$); roadblock avoidance was correlated with (log) navigation response time ($r(381) = -0.302$, $p < 0.001$). A similar pattern was observed when choice quality was substituted for softmax inverse temperature.

As reported in the main text, we also found a relationship between 'Impulsivity' factor scores and softmax inverse temperature, but this appeared to be due to weaker memory for the maze structure, as the effect was no longer significant when including memory performance in the regression model (see **Results**). We found no relationship between 'Worry' factor scores and softmax inverse temperature (TOST equivalence test $t_{(383)} = 2.635$, $p = 0.0438$; coef = -0.0152, 90% CI [-0.0996, 0.0691]), indicating that an effect stronger than $d = 0.15$ could be ruled out. Further, for both roadblock avoidance and navigation response time, associations with both 'Worry' and 'Impulsivity' scores were statistically equivalent to zero within bounds of $d = 0.15$ (TOST p-values < 0.027).

Previous research has identified links between a CIT factor and model-based behavior in a two-step decision task (Gillan et al., 2016; Patzelt et al., 2019; Seow et al.,

2021; Sharp et al., 2023). As noted above, our 'Impulsivity' factor correlated with the derived CIT factor from Hopkins et al. (2022) in our sample (r(382) = 0.826; **Figure S4**).

We found no clear relationships between the CIT factor and measures of goal-directed behavior. The derived CIT factor was not related to softmax inverse temperature ($t_{(382)}$ = -1.009, p = 0.314), model-based roadblock avoidance ($t_{(381)}$ = 0.0584, p = 0.954), or navigation response time ($t_{(382)}$ = -0.177, p = 0.860; all uncorrected). Equivalence tests confirmed that effects of d = 0.15 or larger could be ruled out for all three measures (TOST p-values < 0.024).

However, in an exploratory analysis that included the CIT factor alongside approximations of the two remaining Gillan et al. (2016) factors – anxious-depression (AD) and social withdrawal (LSAS total score) – we found that the CIT factor was negatively correlated with softmax inverse temperature ($t_{(379)}$ = -2.681, p = 0.0077, uncorrected), such that higher CIT scores were associated with lower value sensitivity in choice. This association remained significant when controlling for location memory performance. The corresponding association with the model-free choice quality measure was not significant (p = 0.067). Exploratory analyses of roadblock avoidance and navigation response time still showed no relationships with CIT factor scores in the model including three factors (CIT, AD, and LSAS; p-values > 0.18).

For the episodic influence measures, defined as the tendency to choose previously successful over previously failed goals on day two, we found that the derived CIT factor was positively associated with preferential choice of previously successful over previously failed goals ($t_{(374)}$ = 1.971, p = 0.050, uncorrected). However, this effect indicates that higher CIT scores were associated with a stronger episodic influence on next-day choices. This direction is inconsistent with the negative CIT–softmax association, which would predict weaker sensitivity to task-relevant information at higher CIT scores.

Next, for the derived anxious-depression factor (Gillan et al., 2016; Hopkins et al., 2022), we found that the derived AD factor correlated very highly with our 'Apathy-Anhedonia' factor (r(382) = 0.932), while the correlation between the AD factor and our 'Worry' factor was lower (r(382) = 0.545). The derived AD factor was essentially the same as a factor derived using the weights from Wise & Dolan (2020) (r > 0.99).

As expected given its high correlation with 'Apathy-Anhedonia' factor scores, the AD factor showed a similar but weaker pattern of associations with goal-directed behavior. The derived AD factor was positively related to softmax inverse temperature ($t_{(382)}$ = 1.987, p = 0.0476, uncorrected). However, while the derived AD factor showed a positive association with model-based roadblock avoidance ($t_{(381)}$ = 1.951, p = 0.0518, uncorrected) and faster navigation response time ($t_{(382)}$ = -1.703, p = 0.0895, uncorrected), these associations were not significant. Including the AD factor alongside the CIT factor and LSAS scale in the regression model, to approximate the three factors of Gillan et al. (2016), yielded qualitatively similar results and a slightly stronger roadblock association (p = 0.0362, uncorrected).

#### 7. Goal-directed task earnings and Apathy-Anhedonia

We also examined the relationship between 'Apathy-Anhedonia' factor scores and reward earnings in the choice phase, as noted in the **Results** section. Reward earnings were adjusted for individual differences in risk preferences to account for online titration of goal values. On each trial in which a goal was achieved, the reward amount was divided by the distance-specific scale factor to remove the effect of individual titration. The adjusted reward earnings measure thus better reflects composite behavior, including both discrimination of goal values during choice and navigation efficiency. The effects reported below were qualitatively unchanged when using unadjusted reward earnings.

Earnings were correlated with the influence of goal values on choice (softmax inverse temperature or choice quality; r-values > 0.55). Further, we found that 'Apathy-Anhedonia' factor scores were positively associated with choice phase earnings (r(380) = 0.194, p < 0.001). The relationship remained significant when including either softmax inverse temperature or choice quality in the regression model (p-values < 0.007), and also when additionally controlling for variability in experienced step success rate (p-values < 0.006). Thus, the earnings effect was not driven by random variability in step failure rates across participants. The earnings effect provides additional support for a relationship between 'Apathy-Anhedonia' factor scores and enhanced goal-directed behavior. When choice behavior was accounted for, the residual earnings effect may reflect more efficient navigation in participants with higher ‘Apathy-Anhedonia’ scores, including fewer turn-around events and wall hits.

#### 8. Goal-directed behavior: null relationship with rumination

One potential interpretation of the positive association between goal-directed behavior is that individuals with higher ‘Apathy-Anhedonia’ scores tend to ruminate more. If rumination is directed in part toward the task structure, this may improve goal-directed behavior. While excessive rumination is often considered to detract from task performance, it is possible that in some conditions it improves performance. Our surveys included one item from the Rumination Reflection Questionnaire (“I tend to ‘ruminate’ or dwell over things that happen to me for a really long time afterward”) which has been found to be the strongest (Hur et al., 2017; Trapnell and Campbell, 1999) or second-strongest (Castro et al., 2022) item contributing to a rumination or repetitive thought factor.

Scores on the rumination item were more strongly correlated with ‘Worry’ ($r(382) = 0.721$, $p < 0.001$) than with ‘Apathy-Anhedonia’ scores ($r(382) = 0.471$, $p < 0.001$). In exploratory analyses, we found that scores on the rumination item were not significantly related to softmax inverse temperature ($r(382) = 0.0373$, $p = 0.467$, uncorrected), roadblock avoidance ($r(381) = 0.0313$, $p = 0.541$, uncorrected), or navigation response time ($r(382) = -0.0388$, $p = 0.448$). TOST equivalence tests indicated that we can rule out effect sizes of $d = 0.15$ or larger (TOST p-values $< 0.016$). Thus, while our interpretation is limited by the inclusion of only a single representative rumination item, we found no relationships between rumination and enhanced goal-directed behavior.

#### 9. Reward modulation of HDDM drift rate and Apathy-Anhedonia

In supplemental analyses, we tested a process model of value-based decision-making for goals using a hierarchical drift diffusion model (HDDM), which fits goal decisions and decision reaction times in the choice phase (Wiecki et al., 2013). The key parameter of interest was the modulation of drift rate by goal reward value. The modulation of drift rate by reward is conceptually similar to the model-estimated softmax inverse temperature and model-free choice quality measure, reflecting how well choices are guided by offered values. Importantly, the HDDM additionally accounts for variability in choice reaction time.

We found that the modulation of drift rate by goal value was positively correlated with ‘Apathy-Anhedonia’ factor scores ($t_{(380)} = 3.036$, $p = 0.0026$, including three factors; $r(382) = 0.131$, $p = 0.0103$, bivariate model). This indicates that the effect of goal reward value on decisions and reaction times was stronger in participants with higher ‘Apathy-Anhedonia’ scores. As expected, the modulation of drift rate by goal value effect was correlated with softmax inverse temperature ($r(382) = 0.661$, $p < 0.001$). This additional finding aligns with primary findings of a positive correlation between ‘Apathy-Anhedonia’ factor scores and the modulation of choices by values, but provides a more process-level account of how this association may arise.

#### 10. Risk-taking for goals and additional tasks

We tested for relationships between risk-taking for goals and behavior in the additional tasks. We found that risk-taking for goals was positively related to the risk-seeking parameter in the described gain condition ($r(374) = 0.174$, $p < 0.001$) and negatively correlated with the loss aversion parameter in the described mixed condition ($r(374) = -0.159$, $p = 0.002$). This indicates that participants who took more risks in the goal task were more risk-seeking overall for described prospects. Behavioral measures derived from the delay discounting, effort-reward, and reward learning tasks were unrelated to risk-taking for goals (p-values > 0.20).

For subjective goal success ratings, in contrast to choice behavior, no relationships with additional task variables were observed.

Overall, additional related tasks, specifically decisions for described prospects, explained relatively little variance in risk-taking for goals. Moreover, as the described risk task immediately followed the risky goal task, performance may have been influenced by participants’ extensive prior engagement with risk-related decisions. In general, this suggests that the risk-taking for goals task captures unique features of decision-making behavior.

#### 11. Additional tasks and transdiagnostic factors

In this section, we report relationships between transdiagnostic factor scores and choice behavior in the additional tasks. As decisions in the risky goal task involve features that are also present in common decision-making and learning tasks, we collected data from four additional tasks: a described risk task (gain and mixed conditions), delay discounting, reward learning, and effort-reward decision-making.

First, we investigated the specificity of the novel positive relationships observed between 'Apathy-Anhedonia' factor scores and goal-directed behavior measures. When including all three transdiagnostic factors in the model, we found no associations in the additional tasks between 'Apathy-Anhedonia' factor scores and softmax inverse temperature or the analogous choice quality measure (**Table S4**). Similarly, the positive association with earnings in the risky goal task did not extend to an equivalent measure (summed choice expected value of chosen options) in either condition of the described risk task (p-values > 0.66).

We hypothesized that there would be no association between risk preferences in the described risk task or time preferences in the delay discounting task and depression-related symptoms. Consistent with this hypothesis, we found no significant relationships with 'Apathy-Anhedonia' factor scores (p-values > 0.45; **Table S4**). In exploratory analyses, we found no significant relationships between risk or time preferences and other factors except for a positive association between delay discounting (log(k)) and 'Impulsivity' factor scores ($r(371) = 0.167$, $p = 0.002$, uncorrected; **Table S4**).

We then explored relationships between all three transdiagnostic factor scores and behavior in the remaining two tasks. In the reward learning and effort-reward tasks, we found no significant associations between task behavior and factor scores at an uncorrected level (**Table S4**).

#### 12. Mood at baseline

Baseline mood was computed as the average of happiness ratings collected before each task phase. Across day one and day two, baseline mood was stable across days ($r(384) = 0.775$, $p < 0.001$) but showed meaningful individual variability. We hypothesized that participants with higher depression-related symptoms would report lower mood during the experiment (**hypothesis 7**). Average mood was negatively correlated with all three factors in bivariate analyses (**Figure S10**). However, only the 'Apathy-Anhedonia' factor showed a significant correlation with mood when controlling for the other two factors (day one: $t_{(380)} = -6.921$, $p < 0.001$; day two: $t_{(380)} = -8.392$, $p < 0.001$). Although the other two factors had numerically negative associations with baseline mood, these effects were not significant (day one corrected p-values > 0.05; day two corrected p-values > 0.35).

For individual scales, all were negatively correlated with mood, except the PVSS, which was positively correlated with mood, with all correlations significant at an uncorrected level ($p < 0.05$; **Figure S10**). The anhedonia scale (MASQ-AD) exhibited the strongest negative correlations with baseline mood, followed by the SDS.

#### 13. Emotional responses to outcomes

We hypothesized that participants with higher depression-related symptoms would have blunted emotional responses to goal outcomes (success and failure; **hypothesis 9**). Emotional responses were measured via occasional post-outcome happiness ratings (Rutledge et al., 2014). Overall, post-outcome ratings in the choice phase on day two showed a significant linear effect of outcome (goal 65.7 ± 18.9% [SD]; certain 61.3 ± 19.8%; failure 55.0 ± 21.7%; $t = 12.58$, $p < 0.001$, multilevel model; **Figure S14**).

'Apathy-Anhedonia' factor scores were not related to variability in emotional responses to goal versus certain outcomes ($p > 0.99$, corrected for two comparisons), but scores were associated with blunted responses to certain versus failure outcomes ($r(380) = -0.122$, $p = 0.035$, corrected), supporting our hypothesis.

Exploring the other factors, 'Worry' scores were not significantly related to goal versus certain outcomes ($p > 0.733$, uncorrected) but were related to a larger decrease in response to certain versus failure outcomes ($r(380) = 0.114$, $p = 0.025$, uncorrected). 'Impulsivity' factor scores were not significantly related to goal versus certain outcomes ($p > 0.114$, uncorrected) but were numerically related to blunted certain versus fail outcomes ($r(380) = -0.099$, $p = 0.0541$, uncorrected).

As noted in the **Results** section, including differential emotional response to outcomes as a covariate did not qualitatively change any of the main results.

#### 14. Additional preregistered hypotheses

##### Memory and risk-taking for goals

We hypothesized that weaker memory for the mazes would be related to lower risk-taking for goals (**hypothesis 2**), given that weak memory may increase uncertainty about reaching a goal. Instead, we found that lower memory performance was related to higher risk-taking. Specifically, location memory was negatively related to risk-taking for goals (model-estimated step success probability; day two early: $r(382) = -0.170$, $p < 0.001$; day two end: $r(382) = -0.202$, $p < 0.001$). Additionally, we found a negative association between first-move direction accuracy in the choice phase and risk-taking for goals ($r_s(382) = -0.297$, $p < 0.001$), with a similar effect for the post-test measure of direction accuracy ($r_s(382) = -0.203$, $p < 0.001$). These effects were also observed using a complementary logistic regression-based measure of risk-taking for goals.

The negative relationship between memory measures and risk-taking for goals remained significant when controlling for factor scores, including 'Impulsivity,' which was also negatively related to memory performance.

##### Increase in decision quality across days

We hypothesized that participants' choices would be more guided by presented values with greater experience (**hypothesis 3**). Indeed, we found higher choice quality on day two than day one (day one mean 0.838 ± 0.480 [SD]; day two mean 1.080 ± 0.363; $t_{(402)} = 11.471$, $p < 0.001$).

##### Previous outcome effects

Results regarding **hypothesis 5a–b** and **hypothesis 6a–b**, on the influence of the immediately preceding outcome on subsequent choice, will be the focus of a subsequent report. We do not anticipate additional symptom-related findings beyond those already reported for the relationship between 'Apathy-Anhedonia' and the influence of specific previous episodes on day two choices (**hypothesis 13**).

#### Described prospects risk task and delay discounting task behavior

In this section we report analyses of risk-taking in the described prospects task and the delay discounting task (**hypotheses 15–18**). As noted above, these tasks immediately followed the risky goal task, and thus we cannot determine whether behavior in these tasks was influenced by experience in the preceding risky goal task.

We hypothesized that participants would show risk aversion in the separate described risk tasks (gain and mixed prospects; **hypothesis 15**). In the mixed condition, where the risky option offered a potential gain or loss against a certain option worth zero, we found significant loss aversion (loss aversion parameter mean = 2.64 ± 1.90 [SD], where values above 1 indicate loss aversion; $t_{(387)} = 17.316$, $p < 0.001$), reflected in a low overall risky-prospect choice rate (31.33 ± 23.22%). However, in the gain condition, where the risky option offered a potential gain or zero against a certain gain option, we found risk-seeking (risk parameter = 1.18 ± 1.18, where values above 1 indicate risk-seeking; $t_{(389)} = 8.587$, $p < 0.001$), reflected in a high overall risky-prospect choice rate (78.11 ± 19.55%).

We hypothesized that we would observe significant delay discounting in the separate delay discounting task (**hypothesis 16**) and this was indeed the case (mean k = 0.609, median = 0.0148, where values close to 0 represent minimal discounting; $t_{(374)} = 4.000$, $p < 0.001$).

We hypothesized that these additional tasks would not relate to risk-taking propensity measures from surveys (**hypothesis 17**). This was confirmed in the discounting task, where we found no significant relationship between delay discounting (log(k)) and risk-taking propensity (GRQ general $r(373) = 0.057$, $p = 0.271$; GRIPS $r(373) = 0.060$, $p = 0.244$; **Table S5**).

For described prospects in the gain condition, we found no significant correlation with model-derived risk-seeking (GRQ general $r(374) = 0.0980$, $p = 0.0576$; GRIPS $r(374) = 0.0686$, $p = 0.184$; **Table S5**). However, for the model-free risky gain choice rate, there were significant positive associations with propensity measures (GRQ general $r_s(374) = 0.1230$, $p = 0.0175$; GRIPS $r_s(374) = 0.112$, $p = 0.0302$; all uncorrected).

For described risk prospects in the mixed condition, we found relatively strong negative correlations between the loss aversion parameter and propensity measures (GRQ general $r(374) = -0.244$, $p < 0.001$; GRIPS $r(374) = -0.288$, $p < 0.001$; **Table S5**) and positive correlations between the risky mixed choice rate and propensity measures (GRQ general $r_s(374) = 0.268$, $p < 0.001$; GRIPS $r_s(374) = 0.306$, $p < 0.001$), such that greater choice of mixed prospects (lower loss aversion) was related to higher propensity scores. Regarding the GRQ, correlations were slightly stronger when using the summed GRQ score across all six items rather than the single general item.

We hypothesized that risk-taking would be higher in the risk-taking for goals task than in the described risk prospects task (**hypothesis 18**). This was supported by comparing choice rates for risky options whose expected values were close to the certain alternative, averaging the gain and mixed prospect conditions (goal equal EV choice rate 65.00% versus described prospects 44.09%; $t_{(381)} = 12.549$, $p < 0.001$). However, this pattern was driven by low risk-taking in the described mixed condition (mixed equal EV choice rate 12.65%; gain rate 75.52%). When comparing only with the described gain condition, risk-taking was lower in the risky goals task ($t_{(389)} = -4.362$, $p < 0.001$). As noted above, however, we cannot determine whether behavior in the described risk task was influenced by the immediately preceding risky goal task.

#### Working memory relationships

We hypothesized that performance on the working memory task (modified symmetry-span with letter recall) would be associated with memory performance in the risky goal task (**hypothesis 21**). Working memory was correlated with location memory performance for all measures (early day two: $r_s(358) = 0.188$, $p < 0.001$; all other $r_s$-values > 0.188, p-values < 0.001) and with direction accuracy for all measures (choices on day two: $r_s(358) = 0.153$, $p = 0.0037$; all other $r_s$-values > 0.160, p-values < 0.003). The primary associations between ‘Apathy-Anhedonia’ and goal-directed behavior remained significant when controlling for working memory.

Working memory was positively associated with softmax inverse temperature ($r(358) = 0.139$, $p = 0.0084$, uncorrected), but was not significantly related to model-derived success probability ($p > 0.52$), subjective success ratings ($p > 0.68$), roadblock avoidance ($p > 0.12$), navigation response time ($p > 0.89$), or the influence of previous episodes on day two choices ($p > 0.51$).

Separately, we hypothesized that working memory capacity would be negatively related to risk-taking in the described risk task and to discounting in the delay discounting task (**hypothesis 22**). In the described risk task, we found a negative relationship between working memory performance and loss aversion in the mixed condition ($r(351) = -0.127$, $p = 0.017$), such that greater risk-taking for mixed prospects was associated with higher working memory capacity. We found no relationship with risk seeking in the gain condition ($p > 0.87$). There was a negative but non-significant relationship between working memory performance and delay discounting ($r(349) = -0.087$, $p = 0.104$), such that greater preference for delayed rewards was numerically associated with higher working memory capacity.

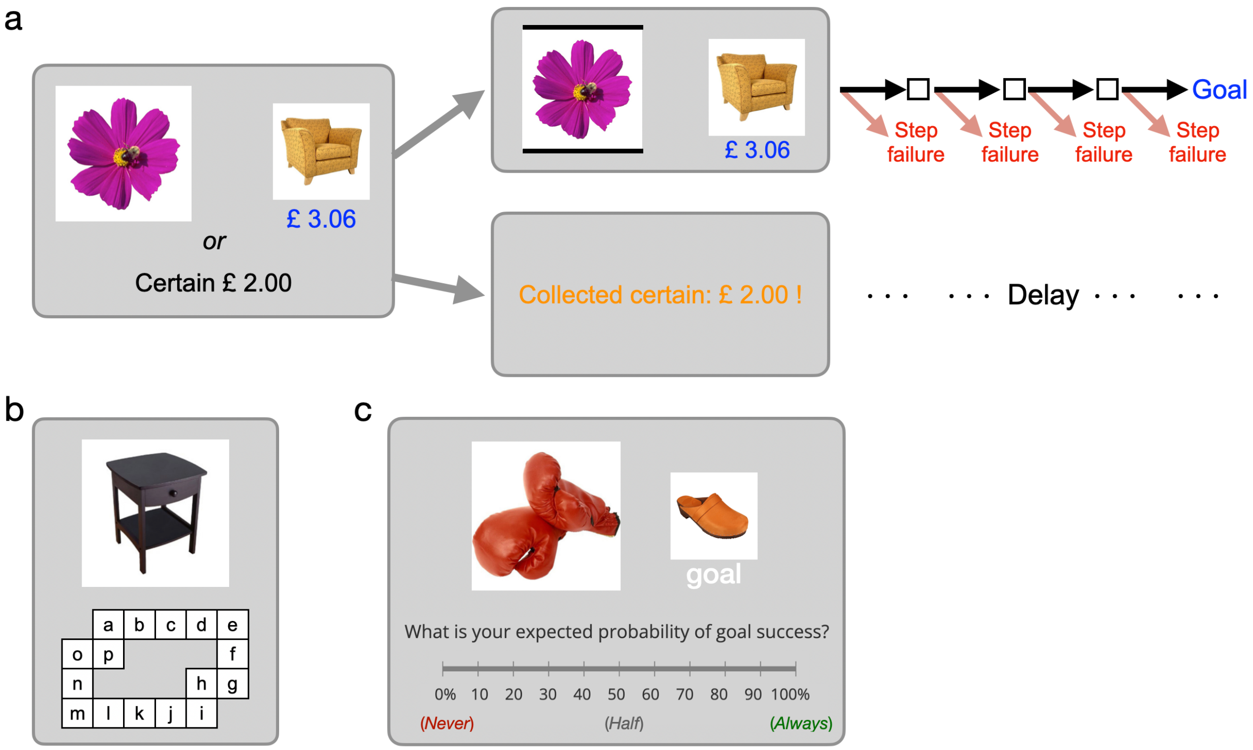


**Figure S1**. Risky goal task trial events, memory test, and success rating displays. a) Sequence of events on a choice trial (see also **Figure 1a-b**). After selecting the goal option, participants navigated using arrow keys. Most moves transitioned to the next location, but each step carried a small probability of a step failure event which terminated the trial. If the certain alternative was selected, the reward amount was displayed, followed by a delay approximating the expected duration of goal navigation. b) Object location replacement memory test. Participants indicated where the displayed object was located in the maze by pressing the corresponding letter key. The outline of the corresponding maze was shown for each object. c) Subjective goal success rating. For specific start–goal combinations, participants rated the probability of successfully reaching the goal while taking the possibility of step failure into account. Ratings were entered via cursor movement on a continuous 0–100% scale.

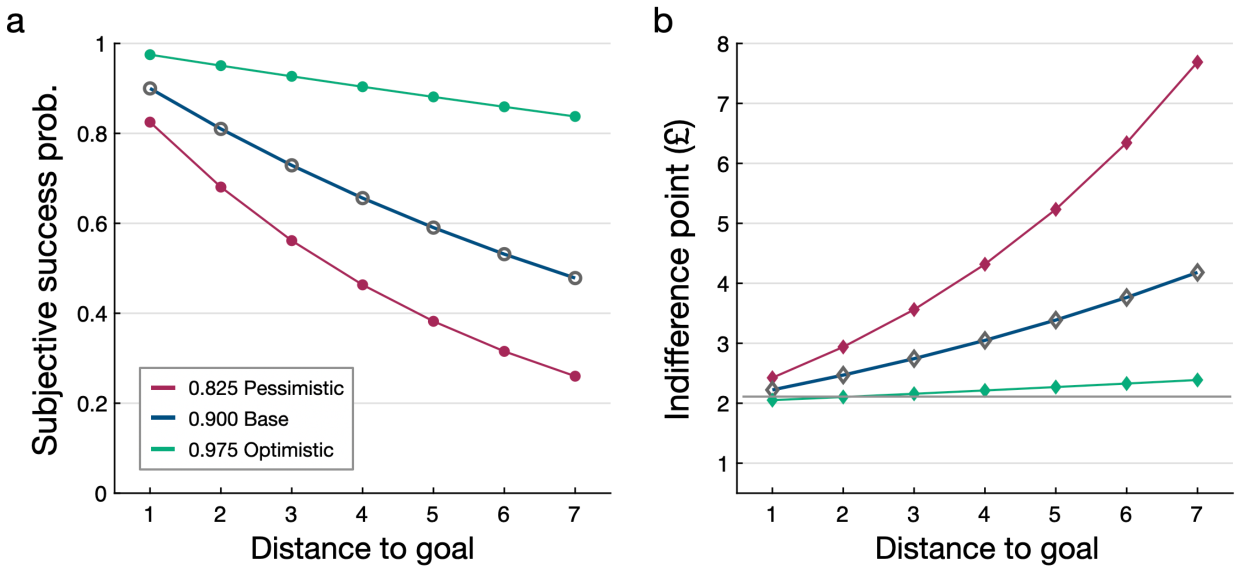


**Figure S2.** Predicted effects of subjective step success rate on goal success probability and indifference points. a) Compounded probability of goal success as a function of distance for three example per-step success rates, illustrating how small differences in per-step success rate are magnified across sequential steps. See **Figure S7** for observed success ratings. b) Predicted indifference points as a function of distance for the same example step success rates. Indifference points are the goal values at which a participant would be equally likely to choose the goal or the certain alternative, . At each indifference point, the probability-discounted goal value equals the £2.00 certain offer. Goal values above the indifference point favor choosing the goal, whereas values below favor the certain option. Lower subjective step success rates produce steeper probability discounting and correspondingly higher indifference points. See **Figure 4** for observed choice behavior.

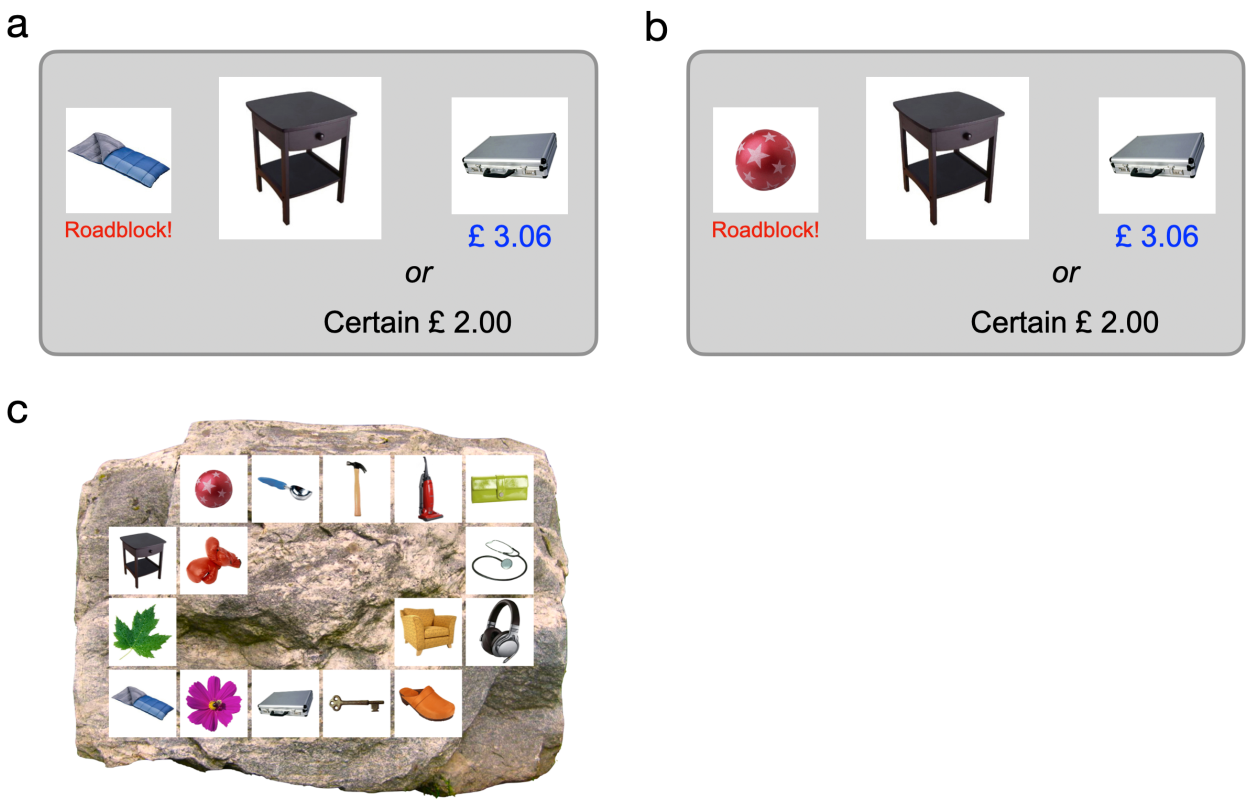


**Figure S3**. Roadblock trial choice screens. a) Relevant roadblock trial: the roadblock is located on the shorter path to the goal, requiring consideration of the longer alternative route. b) Irrelevant roadblock trial: the roadblock is located on the longer path, leaving the shorter path available. c) Maze overview for illustration; this display was not shown during trials.

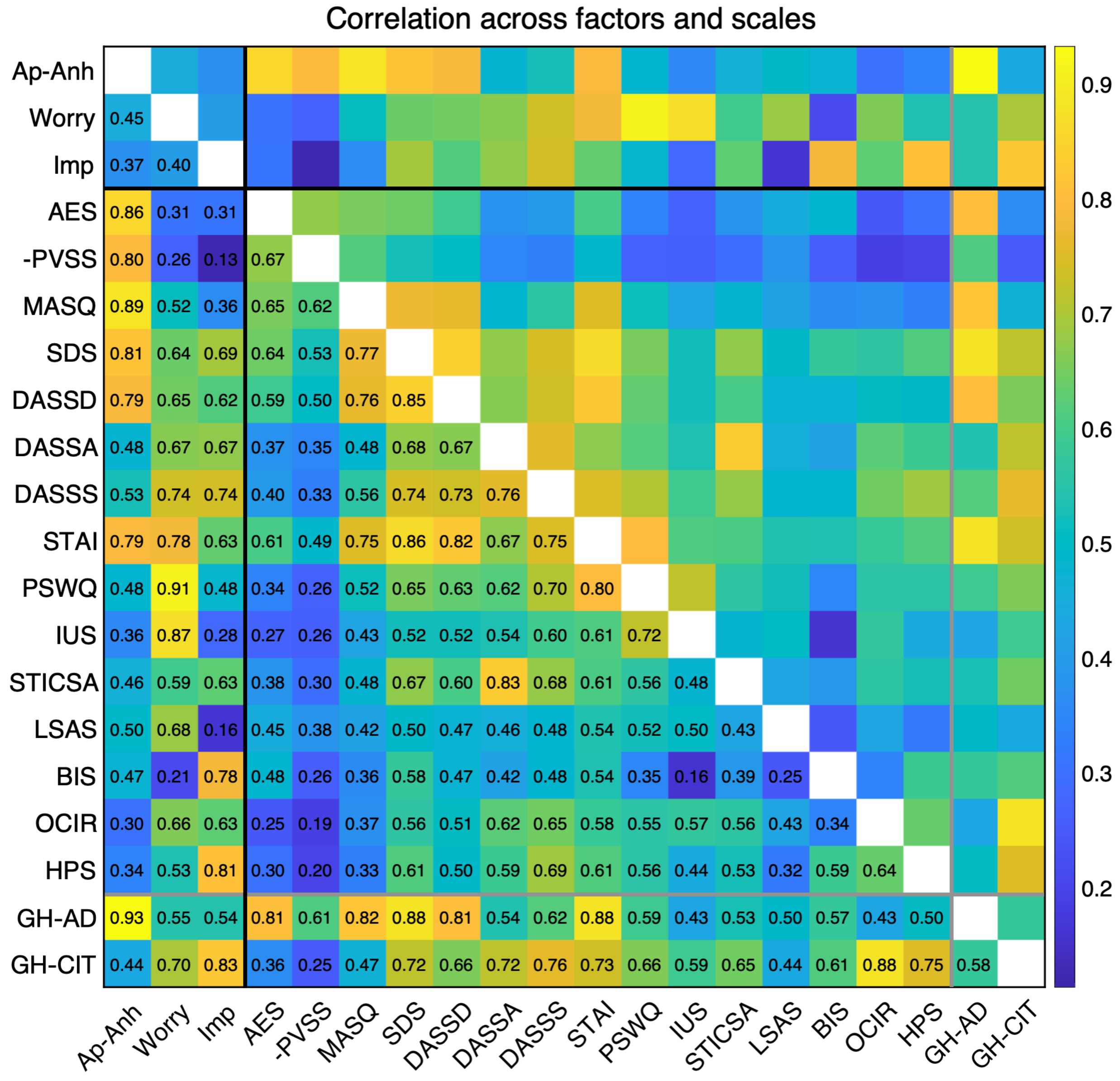


**Figure S4**. Correlations among transdiagnostic factors and individual scales. Pairwise Pearson correlations are shown as a heatmap with correlation values superimposed. PVSS scores were inverted so that higher values indicate greater symptom severity, consistent with the other scales. The DASS-21 was separated into depression, anxiety, and stress subscales. The first three rows and columns, demarcated by black lines, show the three factors identified in the current study (Apathy-Anhedonia, Worry,

Impulsivity). The final two entries, demarcated by gray lines, show derived factors based on the anxious-depression (GH-AD) and compulsivity and intrusive thoughts (GH-CIT) dimensions identified by Gillan et al. (2016), computed using item weights from Hopkins et al. (2022).

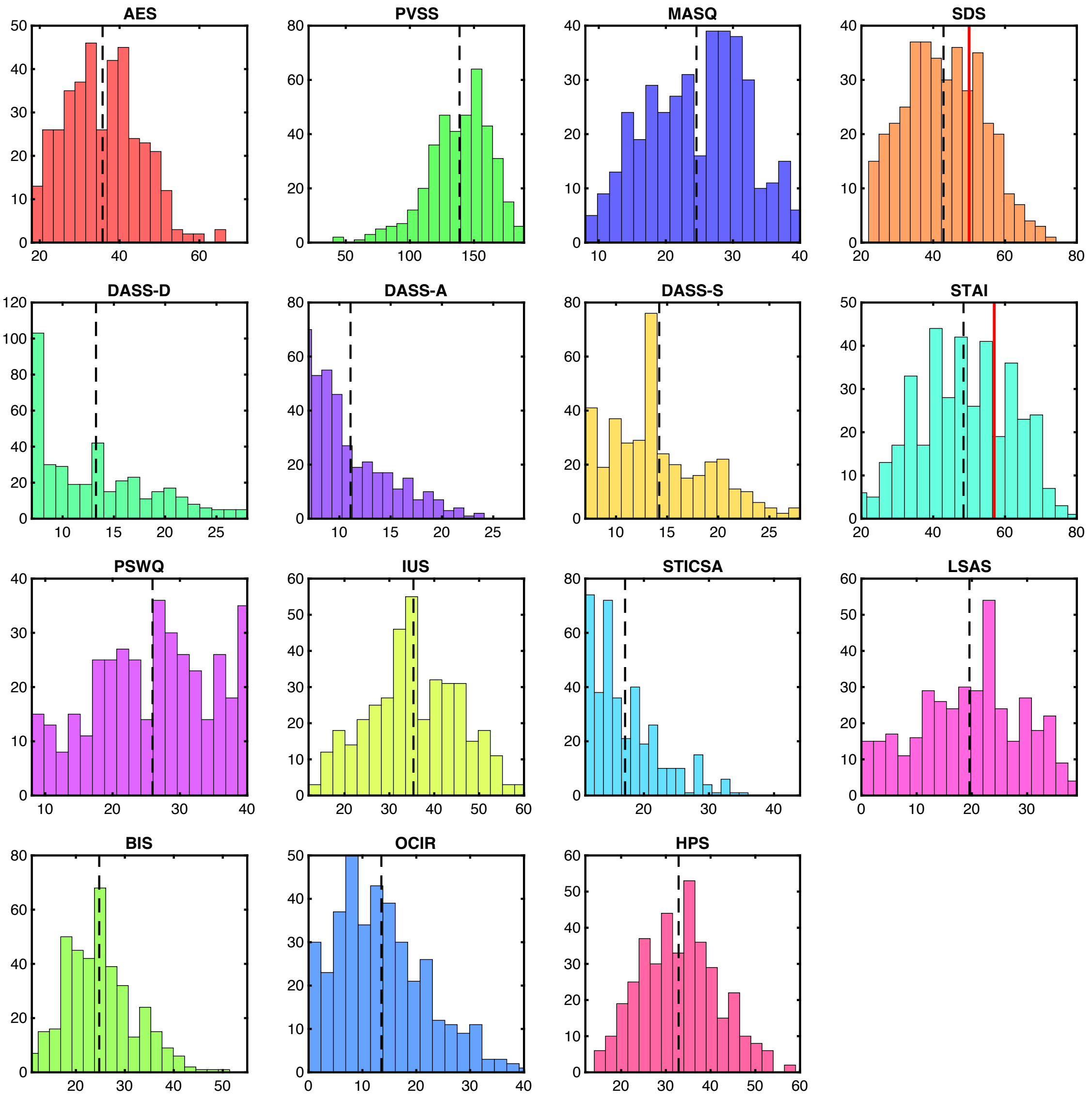


**Figure S5**. Distribution of individual scale scores. Histograms show summed scores for each scale across participants included in behavioral analyses (n = 384). Dotted vertical lines indicate median scores. For each scale, the horizontal axis spans the minimum to maximum possible score. Solid vertical lines indicate clinical reference values: for the SDS, the threshold marks the suggested cutoff for highly symptomatic individuals (>50; Dunstan and Scott, 2019); for the STAI, the threshold marks the reported mean score of a clinical sample with generalized anxiety disorder (> 57; Fisher and Durham, 1999). Only the fear/anxiety ratings were collected for the LSAS, omitting avoidance ratings, because this scale was not of primary interest.

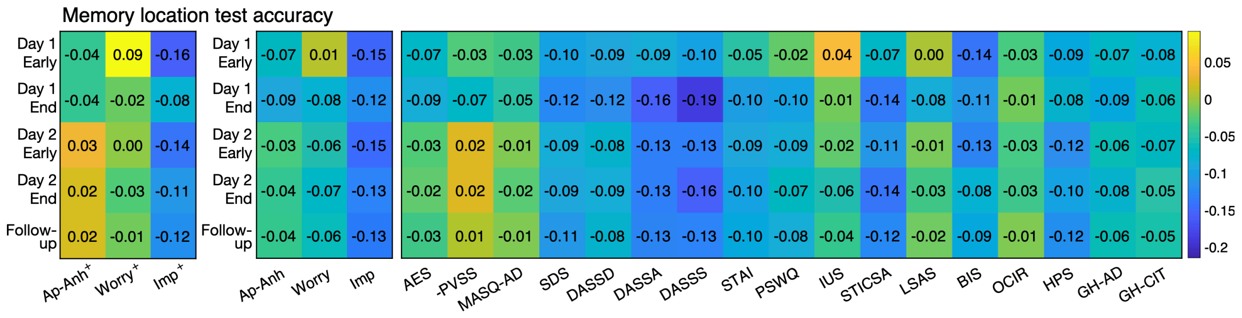


**Figure S6**. Associations between location memory accuracy and symptom measures. Correlation matrix showing the relationship between location memory replacement accuracy (rows: five time points across days) and symptom measures (columns). Left panel: partial Spearman correlations from a regression model including all three transdiagnostic factors simultaneously (denoted by + in the column labels), equivalent to **Figure 3b**. Right panel: bivariate Spearman correlations with the individual transdiagnostic factors and scales. PVSS scores were inverted so that higher values indicate greater symptom severity, consistent with the other scales. Correlation values are superimposed on each cell; correlations exceeding approximately ±0.100 are significant at $p < 0.05$, uncorrected. Sample sizes: day one early, n = 383; day one end, day two early, and day two end, n = 384; follow-up day three, n = 374.

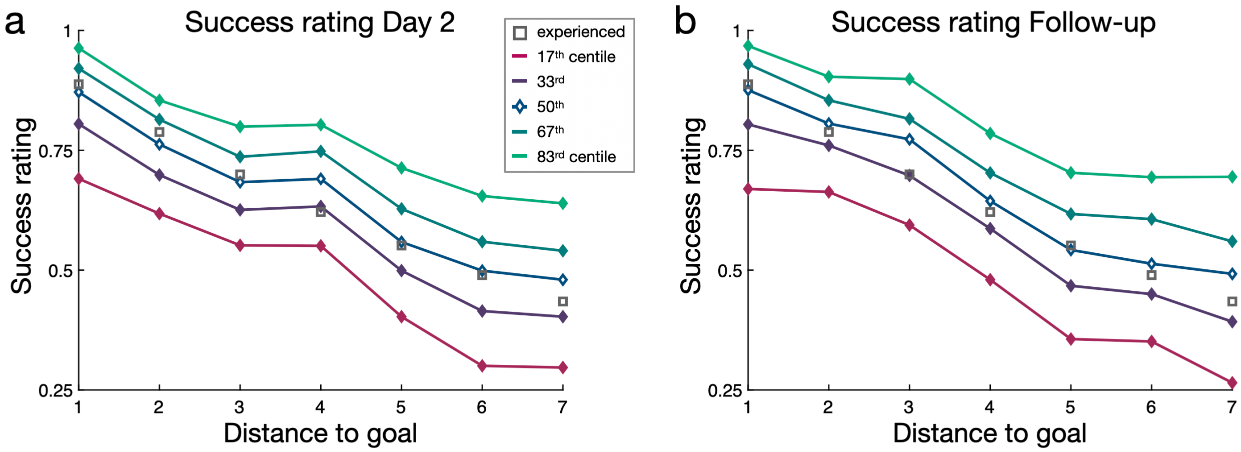


**Figure S7**. Subjective goal success ratings binned by percentile. a) Ratings at the end of day two. b) Ratings at the day three follow-up session. Mean ratings were highly consistent across sessions ($r(373) = 0.810$, $p < 0.001$), and the decrease in success ratings with distance was similar on day two ($t = -31.619$, $p < 0.001$, multilevel model) and day three ($t = -31.453$, $p < 0.001$). Distance was not explicitly provided during success ratings trials; participants were shown only the start image and goal image.

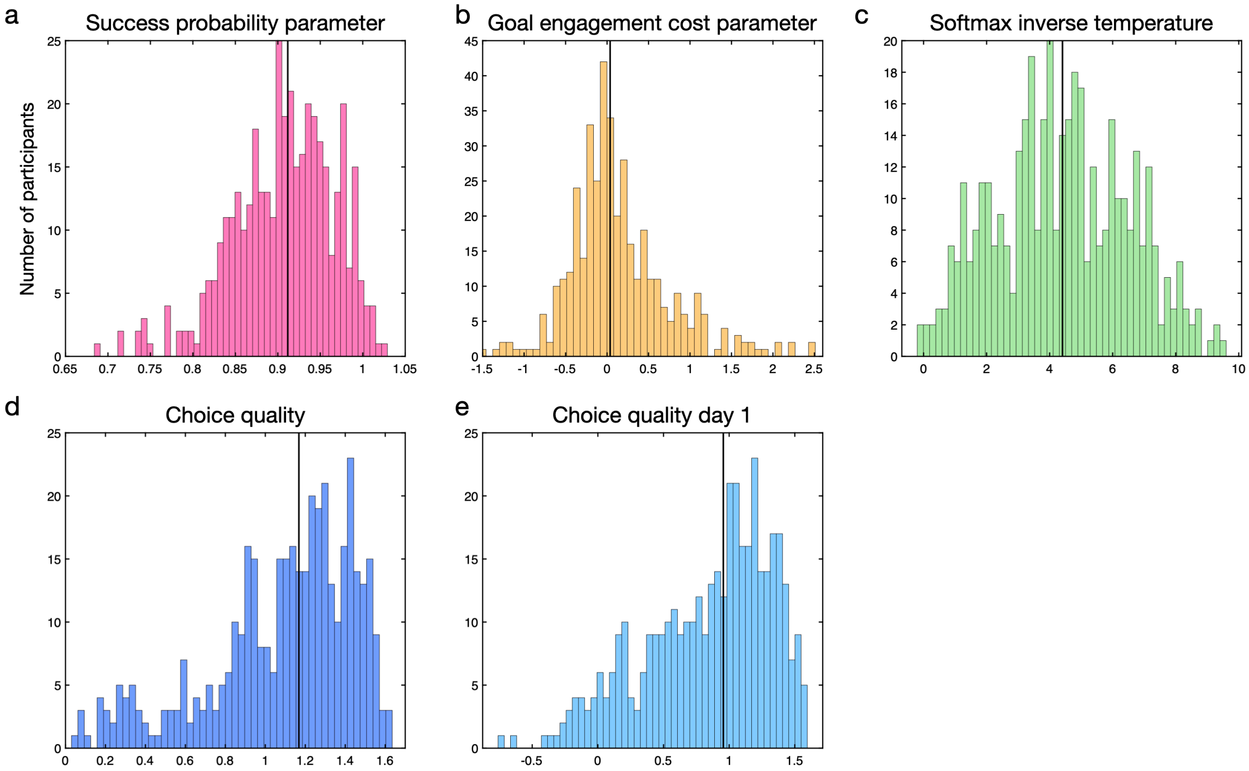


**Figure S8**. Distributions of computational model parameters from the risky goal task and choice quality. a) Per-step success probability parameter. b) Goal engagement cost parameter; the group mean was small but significantly greater than zero (mean £0.145 ± 0.681; $t_{(403)} = 4.801$, $p < 0.001$). c) Softmax inverse temperature, indexing the influence of goal value on choice. d) Model-free choice quality measure, a nonparametric alternative to softmax inverse temperature. e) Model-free choice quality on day one, where the limited number of trials precluded computational model fitting.

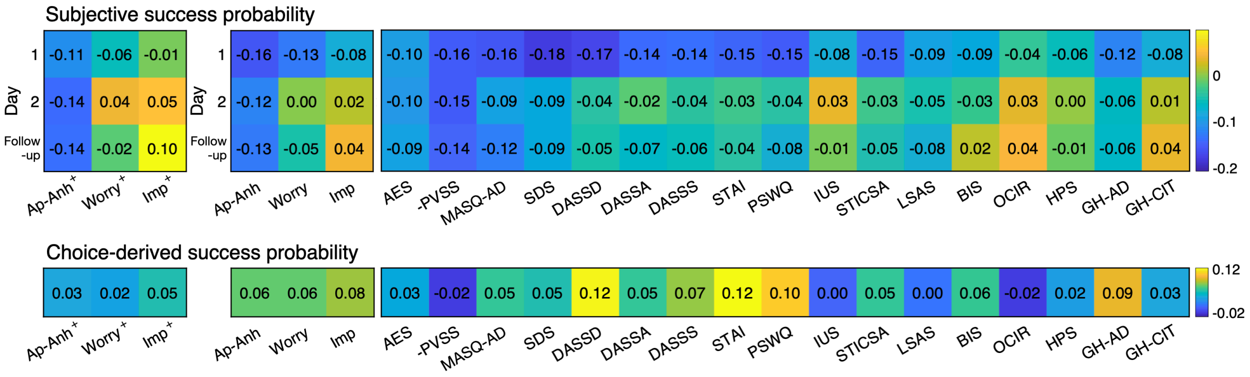


**Figure S9**. Association between belief and choice-derived measures related to goal success and symptom measures. Correlation matrix showing the relationship between subjective success ratings and choice-derived step success probability (rows) and symptom measures (columns). Day two is the primary session for analysis. Left panel: partial Pearson correlations from a regression including all three transdiagnostic factors simultaneously (indicated by + in the column labels), equivalent to **Figure 5**. Right panel: bivariate Pearson correlations with the individual transdiagnostic factors and scales. For ease of interpretation, values of the PVSS scale were inverted. The goal engagement cost parameter is not displayed, as it showed no significant associations with any factor or scale (p-values > 0.11, uncorrected). Correlation values are superimposed on each cell; correlations exceeding approximately ±0.100 are significant at p < 0.05, uncorrected. Sample sizes: subjective success rating day one, n = 382; day two, n = 374; day three, n = 370; choice-derived success probability, n = 380.

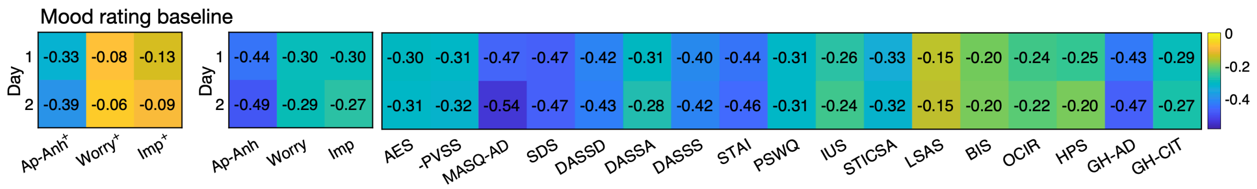


**Figure S10**. Association between baseline mood (happiness ratings) and symptoms measures. Correlation matrix showing the relationship between average baseline happiness ratings (rows: two days) and symptom measures (columns). Left panel: partial Pearson correlations from a regression model including all three transdiagnostic factors simultaneously (denoted by + in the column labels). Right panel: bivariate Pearson correlations with the individual transdiagnostic factors and scales. For ease of interpretation, values of the PVSS scale were inverted. Correlation values are superimposed on each cell; correlations exceeding approximately ±0.100 are significant at $p < 0.05$, uncorrected. Sample sizes: day one and day two, n = 384.

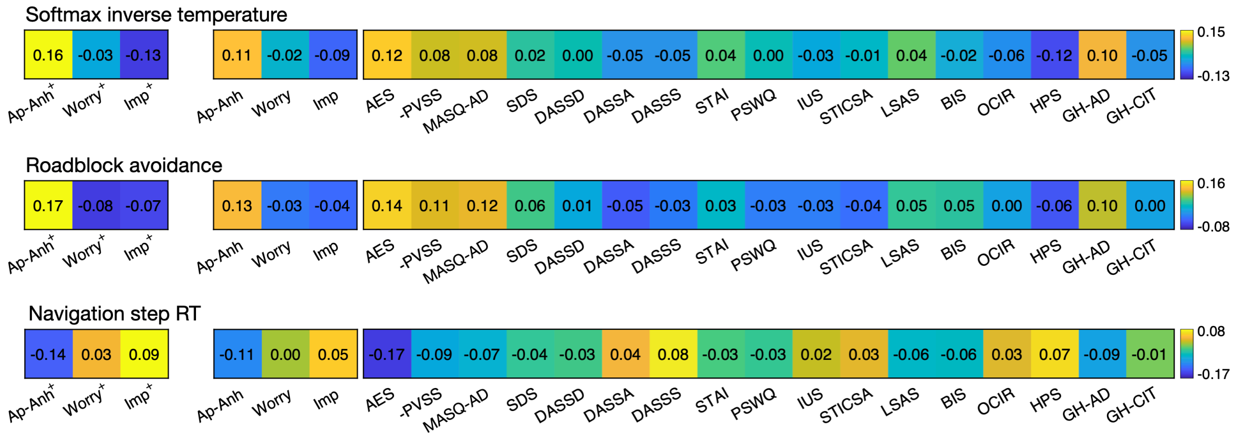


**Figure S11**. Associations between goal-directed behavior measures and symptom measures. Correlation matrix showing the relationship between softmax inverse temperature (sensitivity to goal value), roadblock avoidance, and navigation step RT (rows) and symptom measures (columns). Left panel: partial Pearson correlations from a regression model including all three transdiagnostic factors simultaneously (denoted by + in the column labels), equivalent to **Figure 6**. Right panel: bivariate Pearson correlations with the individual transdiagnostic factors and scales. For ease of interpretation, values of the PVSS scale were inverted. Correlation values are superimposed on each cell; correlations exceeding approximately ±0.100 are significant at $p < 0.05$, uncorrected. Sample sizes: softmax inverse temperature, $n = 384$; roadblock avoidance, $n = 383$; navigation step RT, $n = 384$.

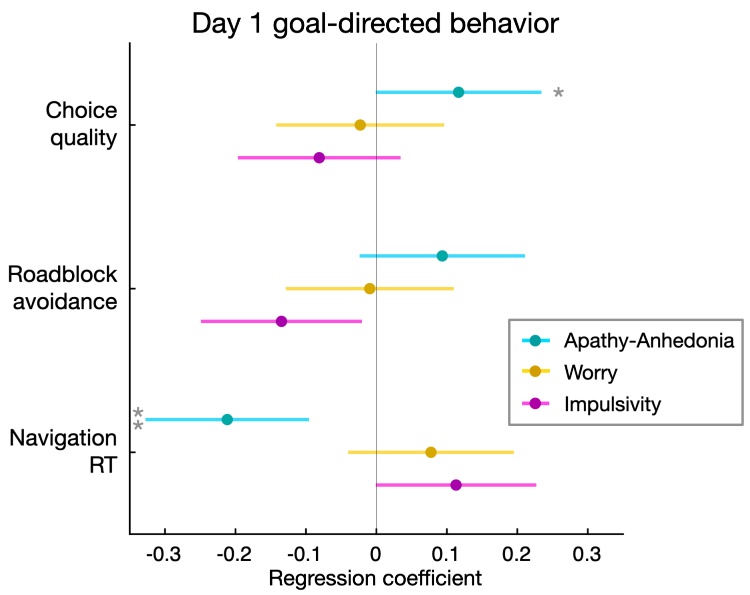


**Figure S12**. Associations between transdiagnostic factor scores and goal-directed behavioral measures on day one, corresponding to the day two analyses in **Figure 6**. Choice quality (indexing the influence of goal value on choice), roadblock avoidance (indexing model-based flexibility), and navigation response time (indexing speed of goal-directed action), are shown. All analyses include the three transdiagnostic factors as predictors. Because the limited number of day one choice trials precluded computational model fitting, the model-free choice quality measure was used. * $p < 0.05$, ** $p < 0.01$.

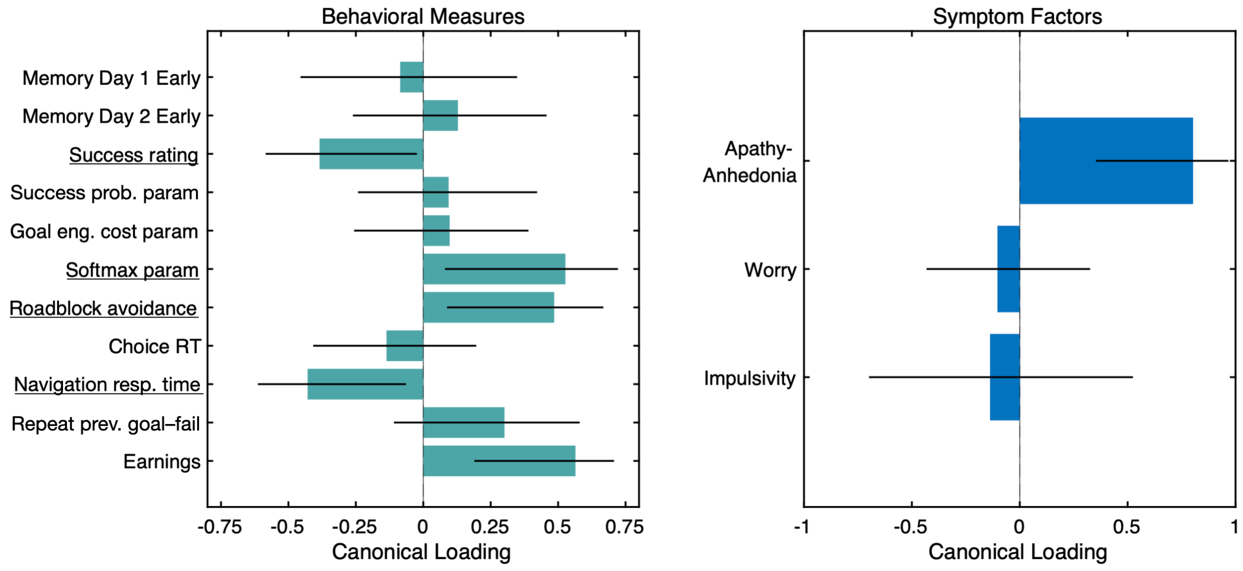


**Figure S13**. Canonical correlation analysis. CCA identified a unified multivariate dimension linking symptom factors to expectations and goal-directed behavior in the first canonical variate pair. Shown are loadings of behavioral measures (left) and symptom factors (right) onto this first canonical variate pair (canonical r = 0.35, permutation p = 0.0003). On the symptom side, 'Apathy-Anhedonia' loads positively, indicating that this dimension drives the multivariate association. On the behavioral side, success ratings load negatively (reflecting pessimistic expectations), while measures of goal-directed performance, including softmax inverse temperature, roadblock avoidance, and faster navigation response time, load in the opposite direction. Error bars denote 95% confidence intervals from a bootstrap procedure (10,000 resamples). The significance of the CCA is determined as a whole; confidence intervals were estimated to illustrate directional consistency rather than the statistical significance of individual variables. Underlined variables on the left indicate primary variables demonstrating significant relationships with 'Apathy-Anhedonia' in the main (non-CCA) analyses. Two secondary variables, choice of previous success versus failure goals and earnings, were also significantly correlated with 'Apathy-Anhedonia' in non-CCA analyses.

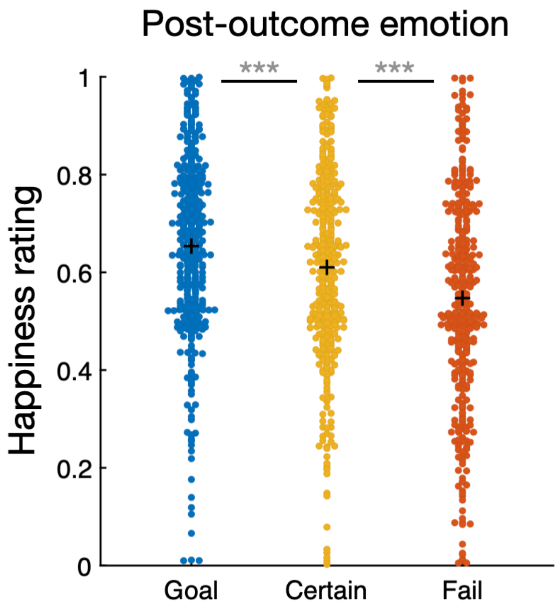


**Figure S14**. Emotional response to choice outcomes during the day two choice phase. Happiness ratings following goal success, step failure, and certain-option choices. *** p < 0.001.

| Model name | IAIC | IBIC | LOO | delta IBIC |
|---|---|---|---|---|
| Exponential discount | 16678.40 | 16716.02 | 16681.58 | 0.00 |
| Hyperbolic | 16758.83 | 16796.45 | 16761.50 | 80.43 |
| Exponential ground truth discount | 17734.53 | 17755.43 | 17734.66 | 1039.41 |
| Exponential without goal engagement cost | 19951.40 | 19972.30 | 19953.69 | 3256.28 |
| Hyperbolic without goal engagement cost | 20982.11 | 21003.01 | 20983.47 | 4286.99 |

**Table S1**. Computational model comparison for risky goal task choices. The exponential discounting model provided the best fit. In both exponential and hyperbolic models, goal value is discounted by cumulative risk, which increases with distance. Model fit was assessed using integrated Akaike Information Criterion (iAIC), integrated Bayesian Information Criterion (iBIC), and leave-one-out cross-validation (LOO). Three control models were also tested: versions of the exponential and hyperbolic models with the goal engagement cost parameter removed, and a “ground truth discount” model in which the per-step success probability was fixed at the targeted 90%. Removing the goal engagement cost parameter substantially worsened fit, as did fixing the success probability.

| Scale Name | Abbreviation | Number of Items | Citation |
|---|---|---|---|
| Apathy Evaluation Scale | AES | 18 | Marin et al., 1991 |
| Positive Valence Systems Scale | PVSS | 21 | Khazanov et al., 2020 |
| Mini Mood and Anxiety Symptoms Questionnaire – Anhedonic Depression | MASQ-AD | 8 | Clark and Watson, 1995 |
| Zung Depression Scale | SDS | 20 | Zung, 1965 |
| Depression Anxiety Stress Scale-21 | DASS-21 | 21 | Lovibond and Lovibond, 1995 |
| State-Trait Anxiety Inventory | STAI | 20 | Spielberger et al., 1983 |
| Penn State Worry Questionnaire | PSWQ | 8* | Kertz et al., 2014 |
| Intolerance of Uncertainty Scale – Short | IUS | 12 | Carleton et al., 2007 |
| State-Trait Inventory of Cognitive & Somatic Anxiety – Somatic Anxiety | STICSA-S | 7* | Ree et al., 2008 |
| Liebowitz Social Anxiety Scale | LSAS | 13* | Liebowitz, 1987 |
| Barratt Impulsivity Scale | BIS | 13* | Patton et al., 1995 |
| Obsessive Compulsive Inventory – Revised | OCIR | 10* | Foa et al., 2002 |
| Hypomanic Personality Scale – Short | HPS | 12 | International Personality Item Pool |

**Table S2**. Summary of mental health questionnaire scales. Scales marked with an asterisk (*) after the item count were not of primary interest and were shortened based on item reduction analyses (Hopkins et al., 2022) and prior psychometric findings. Eight additional individual items were taken from the PHQ-9, FSS, GAD-7, BDI, LOT-R, RRQ, and HADS (two items from the HADS; see **Supplementary Methods** for details).

| Scale | **Mean** | **SD** | **Range** | **% > cutoff** |
|---|---|---|---|---|
| AES | 35.7 | 9.3 | 18 – 65 | |
| PVSS | 138.7 | 24.8 | 45 – 189 | |
| MASQ-AD | 24.6 | 7.4 | 8 – 40 | |
| SDS | 42.9 | 11 | 22 – 73 | 26.2% |
| DASS-D | 13.3 | 5.4 | 7 – 28 | |
| DASS-A | 11.1 | 4.0 | 7 – 24 | |
| DASS-S | 14.2 | 4.8 | 7 – 28 | |
| STAI | 48.5 | 13.1 | 20 – 80 | 27.5% |
| PSWQ (*8 of 16*) | 25.9 | 8.8 | 8 – 40 | |
| IUS-short | 35.3 | 10.1 | 12 – 60 | |
| STICSA-SA | 17.1 | 5.3 | 11 – 36 | |
| LSAS (*13 of 24*) | 19.5 | 9.5 | 0 – 39 | |
| BIS (*13 of 30*) | 24.8 | 7.0 | 11 – 50 | |
| OCIR (*10 of 18*) | 13.5 | 8.4 | 0 – 40 | |
| HPS | 32.9 | 8.6 | 14 – 58 | |

**Table S3**. Descriptive statistics for self-report scale scores in the primary sample (included risky goal task behavior and survey data). For scales with reduced items, numbers in parentheses indicate the number of items included out of the full scale. Clinical reference values are provided for two scales: for the SDS, the threshold marks the suggested cutoff for highly symptomatic individuals (>50; Dunstan and Scott, 2019); for the STAI, the threshold marks the reported mean score of a clinical sample with generalized anxiety disorder (>57; Fisher and Durham, 1999). Only the fear/anxiety ratings of the LSAS were collected, omitting avoidance ratings, as this scale was not of primary interest.

| Task variable | Ap-Anh | Worry | Imp | Ap-Anh + | Worry + | Imp. + |
|---|---|---|---|---|---|---|
| Described gain: risk seeking | 0.02 | 0.10 | 0.10 | -0.04 | 0.07 | 0.07 |
| Described mixed: loss aversion | 0.02 | 0.06 | -0.03 | 0.00 | 0.07 | -0.06 |
| Described gain + mixed: softmax | -0.02 | -0.11* | -0.12* | 0.06 | -0.09 | -0.09 |
| Discounting: log(k) | 0.03 | 0.04 | 0.17** | -0.02 | -0.02 | 0.17** |
| Discounting: softmax | 0.02 | -0.03 | -0.05 | 0.05 | -0.03 | -0.06 |
| Reward learning | 0.03 | 0.00 | -0.08 | 0.05 | 0.02 | -0.10 |
| Effort: effort sensitivity | -0.06 | -0.06 | 0.02 | -0.05 | -0.06 | 0.06 |
| Effort: reward sensitivity | -0.09 | -0.06 | -0.11* | -0.05 | 0.00 | -0.08 |
| Effort: softmax | 0.08 | 0.08 | -0.04 | 0.07 | 0.08 | -0.09 |

**Table S4**. Associations between behavior on additional tasks and transdiagnostic factor scores. Left columns: bivariate correlations. Right columns: partial correlations from regression models including all three factors simultaneously, denoted by (+). Values are Pearson correlations, or Spearman correlations where the behavioral variable distribution was skewed. Softmax parameter values from the discounting model were log-transformed for correlation analyses. Sample sizes: described gain, n = 375; described mixed, n = 354; discounting, n = 375; reward learning, n = 341; effort, n = 350. * $p < 0.05$, uncorrected; ** $p < 0.01$, uncorrected.

| Task variable | GRQ general | GRIPS |
|---|---|---|
| Described gain: risk seeking | 0.10 | 0.07 |
| Described mixed: loss aversion | -0.24** | -0.29** |
| Discounting: log(k) | 0.06 | 0.06 |
| Reward learning | -0.09 | -0.03 |
| Effort: effort sensitivity | 0.07 | 0.03 |
| Effort: reward sensitivity | 0.07 | 0.03 |

**Table S5**. Associations between behavior on additional tasks and self-reported risk-taking propensity. Bivariate correlations. Sample sizes: described gain and described mixed, n = 375; discounting, n = 375; reward learning, n = 341; effort, n = 350. * p < 0.05, uncorrected; *** p < 0.001, uncorrected.
